# G quadruplex DNA facilitates a pervasive path to homologous recombination

**DOI:** 10.64898/2026.06.05.730358

**Authors:** Bhushan L. Thakur, Priyanka Basak, Simran Khurana, Shalu Sharma, Yen-Hsi Lai, Tai Nguyen, Vijayalalitha Ramanarayanan, Xianzhen Zhao, Pedro J. Batista, Travis H. Stracker, Philipp Oberdoerffer

**Author notes:** Co-correspondence to and. contributed equally (Bhushan L. Thakur, Priyanka Basak, Simran Khurana).

## Abstract

Homologous recombination (HR) requires efficient homology search and strand invasion, yet how homologous templates are identified within nuclear space remains unclear. Here, we identify G-quadruplex (G4) DNA structures as pervasive effectors of template strand invasion and uncover the G4 helicase DHX36 as a potent suppressor of this process. DHX36 loss stabilizes G4s, enhances HR, and accelerates repair of replication-associated DNA breaks. G4-mediated HR depends on the non-canonical strand invasion factors RAD51AP1, WDR48, and USP1, and requires G4 motifs within the homologous repair template. DHX36 loss partially restores HR and PARP inhibitor resistance in BRCA1-deficient cells, while promoting aberrant recombination and Alternative Lengthening of Telomeres (ALT). Together, our findings establish dynamic G4 regulation as a key determinant of homology search, genome maintenance, and recombination fidelity.

## Introduction

Homologous recombination (HR) is a fundamental DNA repair process that protects cells from the cytotoxic and mutagenic consequences of DNA double-strand breaks (DSBs) and broken DNA replication forks (*1, 2*). A rate-limiting feature of HR is the search for and invasion of an undamaged, homologous repair template–generally the newly replicated sister chromatid–to ensure accurate repair of the lesion. Homology search requires resection of the broken DNA ends to form a RAD51-coated singe-stranded (ss) DNA nucleoprotein filament, which initiates template strand invasion (*2*). How the RAD51 filament identifies the homologous repair template in three-dimensional nuclear space remains poorly understood. Cohesion-mediated sister chromatid linkage at the DSB site was recently proposed to direct homology search via chromatin loop extrusion (*3, 4*). However, recognition of the homologous template ultimately requires exposure of the complementary ssDNA. In vitro studies indicate that homology sampling is limited to specific regions of the template and enhanced in the presence of an already open, bubble-like repair template configuration (*5, 6*). We thus hypothesize that DNA secondary structures that expose ssDNA may be a central feature of template scanning in HR.

Consistent with this possibility, transcription-associated R-loops have been implicated in HR downstream of end resection (*7*). R-loops accumulate when nascent RNA reanneals with the transiently displaced coding strand to form an RNA:DNA hybrid-containing, three-stranded DNA bubble. However, R-loop formation is generally restricted to actively transcribed loci (*8*), while the need for error-free repair is not. G quadruplexes (G4s) are four-stranded, G-rich alternative DNA structures that, like R-loops, result in the displacement of the opposing DNA strand (*9*). G4s can stabilize R-loops, yet transcription is not a prerequisite for G4 formation (*9-11*). A recent T2T reference genome-based annotation predicts between 400,000 and 2 million G4s in the human genome, resulting in a frequency of one ∼30 bp G4 sequence per every 2-10 kb (*12*). G4-mediated strand opening would, therefore, provide a broadly available means to facilitate homology search.

Precedent for both G4 and R-loop function in genome maintenance comes from the Alternative Lengthening of Telomeres (ALT) pathway, an telomere elongation mechanism essential for the survival of telomerase-deficient cancers (*13*). DNA/RNA G4s and R loops are enriched at telomeric TTAGGG repeats and the associated UUAGGG telomeric repeat-containing RNA (TERRA) (*9, 13*). Both structures are elevated in ALT and have been shown to facilitate break induced replication (BIR), a specialized sub-pathway of HR that mediates telomeric recombination events and G2/M DNA synthesis to extend telomeres (*11, 14, 15*). However, ALT causes aberrant recombination involving non-sister alleles and concomitant extrachromosomal telomeric repeat formation (*16*), underlining a potential threat to genome integrity that may apply to secondary DNA structures beyond telomeres.

Here, we identify the DEAH-box family G4 helicase DHX36 as a potent suppressor of HR. Combining G4-sensitive recombination assays with a comprehensive dissection of DSB repair and the HR-dependent ALT pathway, our work uncovers DNA G4 stabilization as a key determinant of effective, though potentially pervasive, homology search, shaping HR efficiency, cancer genome maintenance, and responses to G4-targeting therapeutics.

### Identification of the G4 helicase DHX36 as a suppressor of HR

In an effort to identify HR-relevant modulators of DNA secondary structures, we took advantage of a focused single-guide RNA (sgRNA) library targeting the majority of known RNA/DNA helicases, which remain poorly explored in this context. In two orthogonal, pooled CRISPR screens, we assessed the impact of gene inactivation on HR efficiency and PARP inhibitor (PARPi) sensitivity. HR efficiency was assessed using U2OS cells carrying the DR-GFP reporter, which is comprised of a full length *GFP* gene interrupted by a target site for the homing endonuclease I-SceI, and a truncated *GFP* donor sequence serving as repair template. A functional GFP gene is restored via HR through gene conversion following Doxycycline (Dox)-inducible DSB formation (*17, 18*). Enrichment or depletion of sgRNAs in GFP^+^ cells was used as a readout for genes that suppress or promote HR, respectively. Six out of 39 RNA/DNA helicases altered HR efficiency, underlining the importance of secondary structure turnover during HR (**Fig. 1A**). To identify HR effectors with clinical potential, we parallelly assessed PAPRi sensitivity in HR-deficient (HR-D) *BRCA1* KO/*TP53* KO RPE1-*hTERT* cells (*19*). PARPi treatment is selectively toxic in HR-D cancers, but resistance via HR restoration remains a major clinical limitation (*20, 21*). Enrichment of sgRNAs in surviving cells is indicative of HR restoration and PARPi resistance upon target protein inactivation. We focused our subsequent efforts on the DEAH-box helicase 36 (DHX36), which was the only helicase that simultaneously suppressed both HR and PARPi sensitivity (**Fig. 1A**).

**Figure 1.**
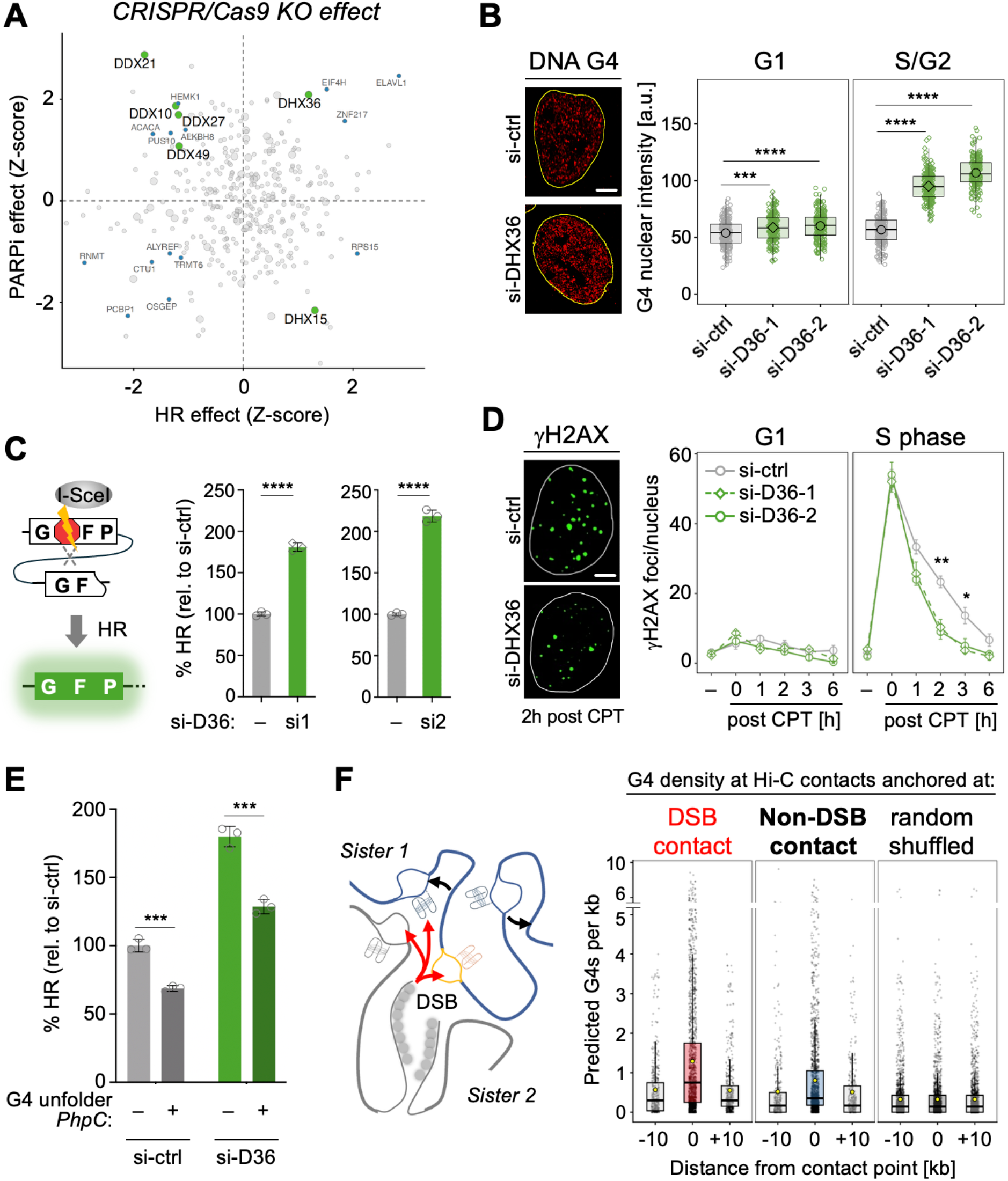
The G4 helicase DHX36 is a suppressor of HR. **(A)** CRISPR/Cas9 screens for modulators of HR and PARPi sensitivity, RNA/DNA helicases are depicted as large circles. Genes meeting cutoff criteria are shown in color (see Methods), helicases are in green. Positive Z scores indicate that genes suppress HR/PARPi resistance. **(B)** DNA G4 abundance in RNAse A-treated U2OS cells expressing the indicated siRNAs, separated by cell cycle stage. Representative S phase nuclei are shown, scale bar: 5 µm, Box plots show mean nuclear G4 intensities for > 250 nuclei per condition. One of two independent experiments is shown. P values are based on the Wilcoxon rank-sum test with BH correction. **(C)** HR efficiency in TRI-DR-GFP U2OS HR reporter cells expressing the indicated siRNAs. Independent DHX36 siRNAs are depicted by distinct symbols, see Fig. S1A. HR efficiency is expressed as the fraction of GFP^+^ cells relative to a control siRNA (–) (n = 3). **(D)** IF analysis of *γ*H2AX foci in U2OS cells expressing the indicated siRNAs. Cells were left untreated or treated with 100 nM CPT for 1h and analyzed at the indicated recovery times in G1 and S phase cells, representative S phase nuclei 2h after CPT are shown, scale bar: 5 µm. Values represent average and S.E.M. of mean foci counts from three independent experiments. P values compare si-ctrl to combined si-DHX36 means. **(E)** HR efficiency as in (B) in TRI-DR-GFP U2OS cells expressing the indicated siRNAs in the presence or absence of 100µM PhcP, symbols represent biological replicates (n = 3 biological replicates). **(F)** Density of predicted G4 forming sequences at Hi-C contacts involving a sgRNA-induced DSB (red arrows), or non-DSB containing Hi-C contacts (black arrows). DSB:DSB contacts are significantly different from flanking regions and non-DSB, p<0.0001 based on two-sided permutation testing. All bar graphs depict mean and S.D., symbols represent biological replicates. Unless indicated otherwise, P values are based on Student’s two-tailed t-test: * P < 0.05; ** P < 0.01; *** P< 0.001; **** P < 0.0001.

### G4 stabilization facilitates HR

DHX36 primarily targets G4 nucleic acid structures and was recently shown to resolve DNA G4s in a DNA replication-dependent manner (*22, 23*), placing DHX36 function in the context of cells that are primed to undergo HR. DHX36 knockdown with two independent siRNAs corroborated increased DNA G4 accumulation in S/G2 phase cells (**Fig. 1B**). We further confirmed the increase in HR observed in our CRISPR KO screen by RNA interference, using at least two independent siRNAs against DHX36 in two distinct cell lines carrying stably inserted DR-GFP knock-in alleles (U2OS cells, **Fig. 1C, Fig. S1A-C** and HEK293T cells, **Fig. S1D**). Cell cycle analyses demonstrate that the effect of DHX36 depletion on HR is not a result of an increase in S/G2 phase cells (**Fig. S1E**). Of note, the DR-GFP transgene contains ∼50 putative G4-forming sequences, spread evenly between the sense and antisense strands (see **Fig. S3B**). G4 accumulation in the reporter locus was confirmed using chromatin immunoprecipitation (ChIP) against G4 DNA, and DHX36 depletion further increased DR-GFP-associated G4 levels, making this a suitable system to study G4 function in HR (**Fig. S1F**). Supporting the latter, treatment with the G4-unfolding compound phenyl-pyrrolocytosine (PhpC) decreased HR efficiency and partially reversed the effect of DHX36 loss, while the G4 stabilizing compound pyridostatin (PDS) had the opposite effect(**Fig. 1E, Fig. S1G**).

Given that HR is essential for the protection and restart of broken replication forks (*1*), we next assessed how DHX36 depletion affects the repair of replication-dependent DSBs. Fork collapse was induced via camptothecin (CPT)-mediated stabilization of Topoisomerase 1 (TOP1) covalent cleavage complexes (TOP1cc), which are converted into single-ended DSBs that rely on HR for efficient repair (*24*). DSBs, detected via immunofluorescence (IF) for histone H2AX phosphorylated on S139 (*γ*H2AX), were readily induced following CPT treatment in an S phase-specific manner in both control and DHX36 knockdown cells. Following release from CPT, cells lacking DHX36 displayed significantly faster clearance of *γ*H2AX foci, consistent with improved HR upon G4 stabilization (**Fig. 1D, Fig. S1H**). Together these findings establish a role for DHX36-regulated G4 turnover in homology-directed DSB repair.

### Inactivation of DHX36 facilitates non-canonical strand invasion

Recent Hi-C mapping of DSB-associated chromatin contacts revealed localized *de novo* DSB-chromatin interactions as a result of DSB induction, reflecting homologous template scanning (*3, 4, 25*). Supporting a possible role for G4 DNA structures in this process, predicted G4-forming sequences were significantly enriched at DSB-containing chromatin contacts compared to contact-distant control regions or non-DSB-associated Hi-C contacts (**Fig. 1F**). We thus conclude that sites where DSBs engage with template DNA often contain DNA sequences with G4 forming potential.

Based on this observation and given that G4 structures can stabilize R-loops, we sought to assess the role of two HR effectors previously implicated in enhancing R-loop-mediated strand invasion: RAD51 Associated Protein 1 (RAD51AP1) and WD40-repeat containing protein 48 (WDR48/UAF1) (*7, 26*). Depletion of either WDR48 or RAD51AP1 abrogated the ability of DHX36 inactivation to promote HR (**Fig. 2A, Fig. S2A-D**). A similar effect was observed for the WDR48/RAD51AP1-interacting protein USP1 (**Fig. S2B**) (*27, 28*). To rule out a defect in RAD51 filament formation, we assessed RAD51 foci in response to ionizing radiation (IR)-induced DNA damage. EdU labeling of S phase cells combined with nuclear size confirmed S/G2 specificity of RAD51 foci (**Fig. S2F**). While DHX36 loss increased overall RAD51 foci numbers through mechanisms that will be explored elsewhere, foci remained elevated upon co-depletion of WDR48 or USP1 (**Fig. S2F**). Moreover, the effect of DHX36 loss was entirely dependent on RAD51 (**Fig. 2B**). DHX36 loss thus promotes HR via non-canonical strand invasion effectors downstream of RAD51 filament formation. Notably, DHX36 loss did not increase RAD51 foci upon co-depletion of RAD51AP1, consistent with a role for RAD51AP1 in RAD51 filament stabilization (*29*) (**Fig. S2F**). In line with the reporter assay, we observed that accelerated repair of CPT-induced, replication-associated DSBs upon DHX36 loss was reverted to control levels following depletion of either RAD51AP1 or WDR48 (**Fig. 2C**).

**Figure 2.**
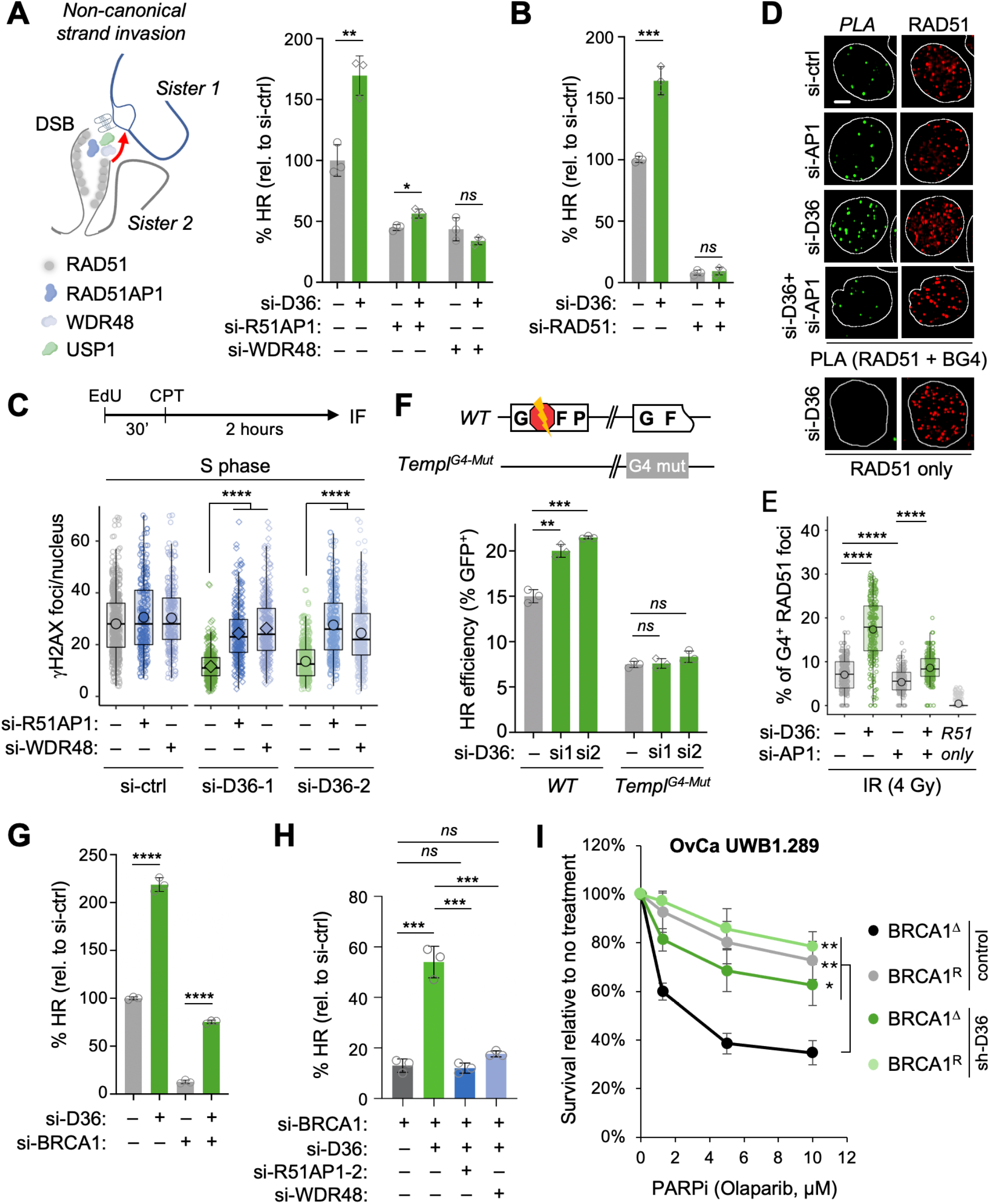
G4 stabilization promotes noncanonical strand invasion. **(A)** Schematic depicting RAD51AP1/WDR48-mediated strand invasion. **(A, B)** HR as in Fig. 1C for the indicated siRNA combinations, “–” indicates addition of control siRNA (si-ctrl) instead of gene-specific siRNA, symbols represent biological replicates (n = 3). **(C)** *γ*H2AX IF following CPT treatment as in Fig. 1D in S phase cells expressing the indicated siRNA combinations. Box plots show mean foci counts from at least 250 nuclei, one of two independent experiments is shown. **(D, E)** Proximity ligation assay between G4 and RAD51 four hours after 4 Gy IR in U2OS cells expressing the indicated combinations of si-DHX36-2 and si-RAD51AP1-2. Representative S phase nuclei are shown for PLA signal and RAD51 foci, scale bar: 5 µm. RAD51 antibody only served as control for non-specific PLA signal (R51 only) (D), box plots depict the fraction of G4-associated RAD51 foci, at least 250 nuclei were analyzed per sample, one of two independent experiments is shown (E). P values in (C) and (E) are based on Wilcoxon rank-sum test with BH correction. **(F)** HR efficiency expressed as % GFP^+^ cells in 293T-DR-GFP and 293T-DR-GFP-Templ^G4Δ^ cells, following transient I-SceI transfection. G4 mutations were restricted to the template region of the reporter, symbols represent biological replicates (n = 3). **(G, H)** HR assays as in (A), symbols represent biological replicates (n = 3). **(I)** Colorimetric survival assay of BRCA1^Δ^ UWB1.289 ovarian cancer cells and isogenic, BRCA1-reconstituted counterparts (BRCA1^R^) expressing shRNA against DHX36 or empty vector (control). Cells were treated with Olaparib at the indicated doses for 72h. Average and S.E.M from four independent experiments are shown. All bar graphs depict mean and S.D., symbols represent biological replicates. Unless noted otherwise, P values are based on Student’s two-tailed t-test. (C, E): * P < 0.05; ** P < 0.01; *** P< 0.001; **** P < 0.0001.

Collectively, our findings thus far are consistent with a role for template strand G4s in directing RAD51AP1/WDR48-mediated homology search. To test this, we assessed the frequency of RAD51 colocalization with G4 DNA using the proximity ligation assay (PLA), which allowed us to quantify IR-induced RAD51 repair foci within 40 nm of a G4 structure in the presence or absence of DHX36. G4:RAD51 colocalization was defined as the fraction of PLA foci relative to the total number of RAD51 foci to account for knockdown-intrinsic variations in RAD51 foci counts. We readily observed IR-induced G4 colocalization in ∼8% of RAD51 foci in wildtype cells, which increased more than two-fold upon DHX36 depletion (**Fig. 2D, E**). Co-depletion of RAD51AP1 significantly reduced DHX36 loss-driven RAD51:G4 association, despite similar nuclear G4 levels (**Fig. 2D, E, Fig. S3A**). We therefore conclude that G4 stabilization upon DHX36 loss promotes RAD51 filament association with G4s in a RAD51AP1-dependent manner.

To formally test whether the presence of G4s specifically in the repair template can facilitate HR, we mutated all predicted G4-forming motifs in the template region of the DR-GFP reporter. G4s flanking the DSB site were left unaltered (**Fig. S3B**). Both the mutant Template^ΔG4^ DR-GFP and wildtype DR-GFP transgenes were inserted into the AAVS1 locus of HEK293T cells (**Fig. S1C**). HR was readily detected in the Template^ΔG4^ reporter, although it was reduced compared to the WT construct, consistent with a minor reduction in sequence homology between the DR-GFP DSB site and template region. Consistent with our findings in U2OS cells, inactivation of DHX36 with two independent siRNAs significantly increased HR in the WT DR-GFP transgene (**Fig. 2F**). In contrast, DHX36 depletion failed to promote HR in the absence of G4 forming sequences in the homologous template sequence (**Fig. 2F**). We note that 293T cells have higher basal HR levels at the DR-GFP locus compared to U2OS cells (Fig. 2F vs. Fig. S1D), which likely accounts for the relatively modest increase in HR following DHX36 inactivation in 293T cells. Together, these findings reveal that G4 stabilization in the repair template promotes HR downstream of RAD51 filament formation via a noncanonical strand invasion pathway.

### G4-mediated HR can partially restore BRCA1 loss

DHX36 depletion not only increases HR efficiency but also protects from PARPi toxicity in BRCA1-deficient cells (**Fig. 1A**). We, thus, sought to interrogate the functional relationship between BRCA1 loss and G4-mediated HR. BRCA1 controls multiple aspects of the HR cascade from end resection and RAD51 filament formation to template strand opening and invasion (*2, 6, 30*). Consistent with this, BRCA1 depletion almost completely abrogated HR in our reporter system (**Fig. 2G, Fig. S2A**). Co-depletion of DHX36 with two independent siRNAs resulted in a significant, 5–10-fold restoration of HR in BRCA1-deficient cells, which was entirely dependent on RAD51AP1 and WDR48 and further involved the restoration of RAD51 filament formation, a prerequisite for strand invasion (**Fig. 2G, H, Fig. S2F**). How DHX36 loss bypasses the need for BRCA1 during RAD51 loading warrants further investigation. Together, these findings establish G4-driven, RAD51AP1/WDR48-mediated recombination as a process that can overcome HR deficiency associated with BRCA1 loss, likely by providing an alternative means for strand invasion. Consistent with this, stable knockdown of DHX36 in BRCA1^Δ^ UWB.289 ovarian cancer cells resulted in PARPi resistance comparable to that observed upon BRCA1 re-expression (**Fig. 1A, Fig. 2I**).

### G4s promote HR independent of transcription

Given that the RAD51AP1 pathway, R-loops, and G4s cooperate in the context of ALT (*11, 15*), we next asked whether G4-mediated HR is dependent on R-loops. Consistent with a role for G4s in R-loop stabilization (*9, 11, 22*), we observed a modest increase in nuclear R-loops following DHX36 depletion, which, like G4 accumulation, was specific to S phase cells (**Fig. 3A**). To assess the contribution of R-loops to HR, we depleted the R-loop resolvase Senataxin (SETX), which resulted in robust R-loop accumulation (**Fig. 3A**) (*31, 32*). In agreement with a previous report (*33*), depletion of SETX increased HR in our reporter system. SETX depletion was epistatic with DHX36 loss (**Fig. 3B**), and analogous to DHX36 depletion, the effect of SETX loss was at least in part dependent on WDR48 and RAD51AP1 (**Fig. 3C, Fig. S4**), indicating that G4s and R-loops cooperate to promote non-canonical strand invasion at DSBs in transcribed genomic regions beyond the telomere.

**Figure 3.**
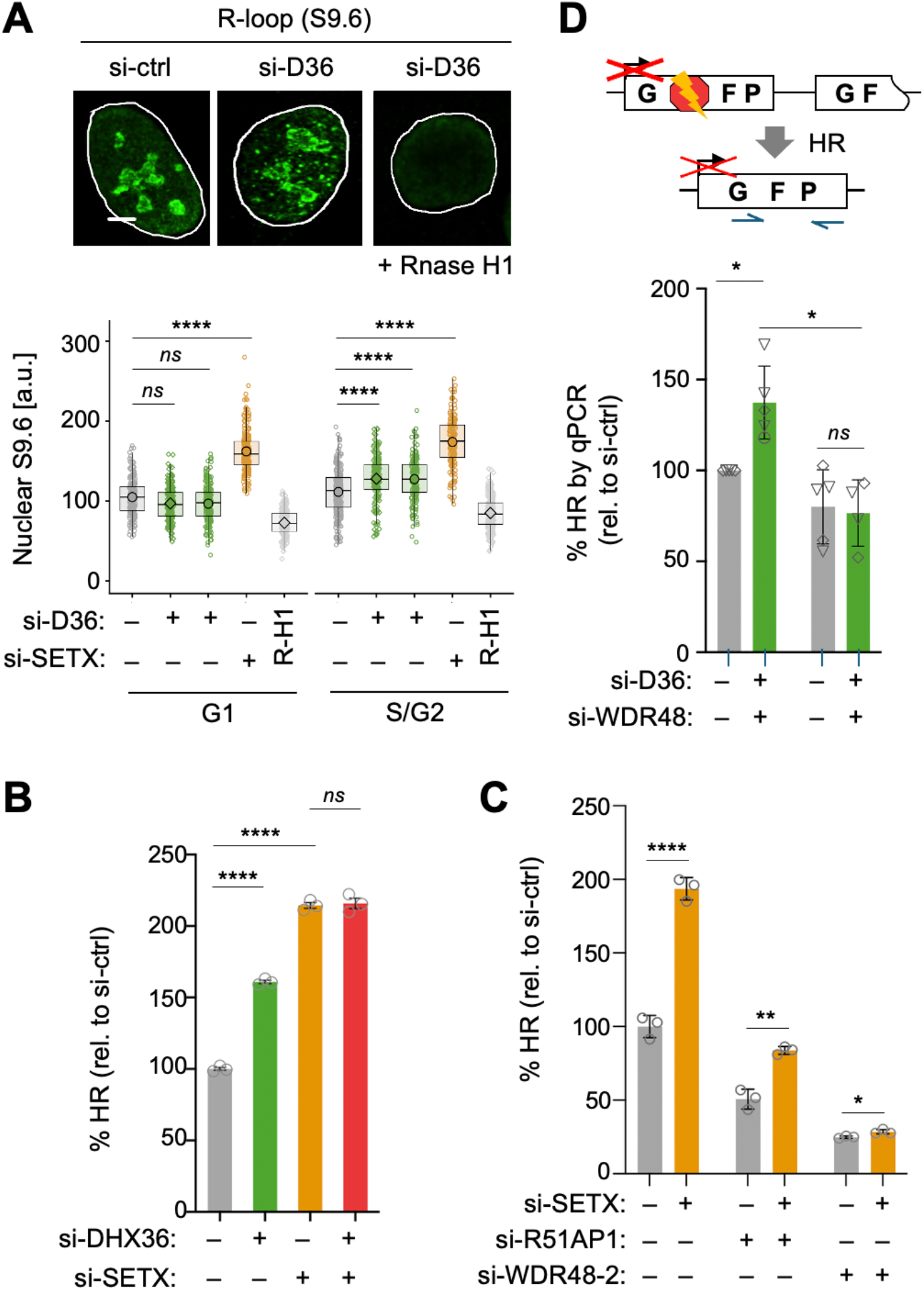
R-loops cooperate with but are not required for G4-mediated HR. (**A)** IF for R-loops (S9.6) in U2OS cells expressing the indicated siRNA combinations in the presence or absence of RNAse H1 in G1 and S/G2 cells. Representative S phase nuclei are shown, scale bar: 5 µm. Box plots show mean intensities from at least 250 nuclei transfected with the indicated siRNA, symbols depict the mean. P values are based on Wilcoxon rank-sum test with BH correction, one of two independent experiments is shown. **(B, C)** HR efficiency in TRI-DR-GFP cells transfected with the indicated siRNA combinations, analyzed as in Fig. 1C, symbols represent biological replicates (n = 3). **(D)** HR assay in cells with a transcriptionally inactive DR-GFP reporter transfected with the indicated siRNA combinations. HR efficiency was assessed by qPCR amplification of the HR repair product, normalized to a control locus. HR is expressed relative to control siRNA. Independent experiments are combined and P values are based on a paired student’s t-test. Independent DHX36 siRNAs are depicted by distinct symbols, see Fig. S1A. All bar graphs depict mean and S.D., symbols represent biological replicates (n ≥ 5). Unless noted otherwise, P values are based on Students two-tailed t-test: * P < 0.05; ** P < 0.01; *** P< 0.001; **** P < 0.0001.

Because G4s can form independently of R-loop formation (*9, 11*), it is conceivable that G4-associated template opening is sufficient to promote HR even in the absence of transcription. To test this, we took advantage of a transcriptionally inactive DR-GFP reporter cell line, in which recombination events are detected via quantitative PCR (qPCR) (*7*). Following DHX36 depletion with two distinct siRNAs, we observed a significant, transcription-independent increase in HR efficiency, which was abrogated in the absence of WDR48 (**Fig. 2G**). We therefore conclude that G4 stabilization upon DHX36 loss can facilitate HR independent of transcriptional activity, positioning G4s as broadly available genomic effectors of strand invasion.

### DHX36 loss triggers aberrant recombination in the context of ALT

While enhancing HR efficiency, pervasive RAD51 contacts with open DNA during homology search carries an inherent risk of aberrant repair and chromosomal aberrations. To assess the potential of DHX36 to suppress aberrant genome maintenance, we analyzed the consequences of DHX36 loss on the ALT pathway (*11, 13, 15*). We examined the effects of DHX36 depletion on ALT using ssTelo-FISH to quantify ssDNA levels as a surrogate for ALT activity (*34, 35*). Depletion of DHX36 led to a marked elevation of ssTelo signal in ALT^+^ U2OS and LM216J cells, however little signal was observed in hTERT^+^ LM216T or HeLa-LT cells (**Fig 4A and Fig S5A, B**). ALT is triggered by the unwinding of lagging strand intermediates by the BLM helicase and/or stochastic strand breakage events that arise due to elevated telomeric repeat-driven replication stress (*13, 36*). Co-depletion of BLM with DHX36 in U2OS cells eliminated the increased signal, supporting the fact that it was a result of canonical ALT activity (**Fig. S5C**). Analysis of both APBs and c-circles further corroborated this result, as each was elevated by DHX36 depletion (**Fig. 4B, C**). Using PLA to measure DNA G4s in proximity to the shelterin component TRF2, we confirmed an increase in telomeric G4s upon DHX36 loss (**Fig. 4D**). These data indicate that DHX36 loss stabilizes telomeric G4s and elevates ALT activity.

**Figure 4.**
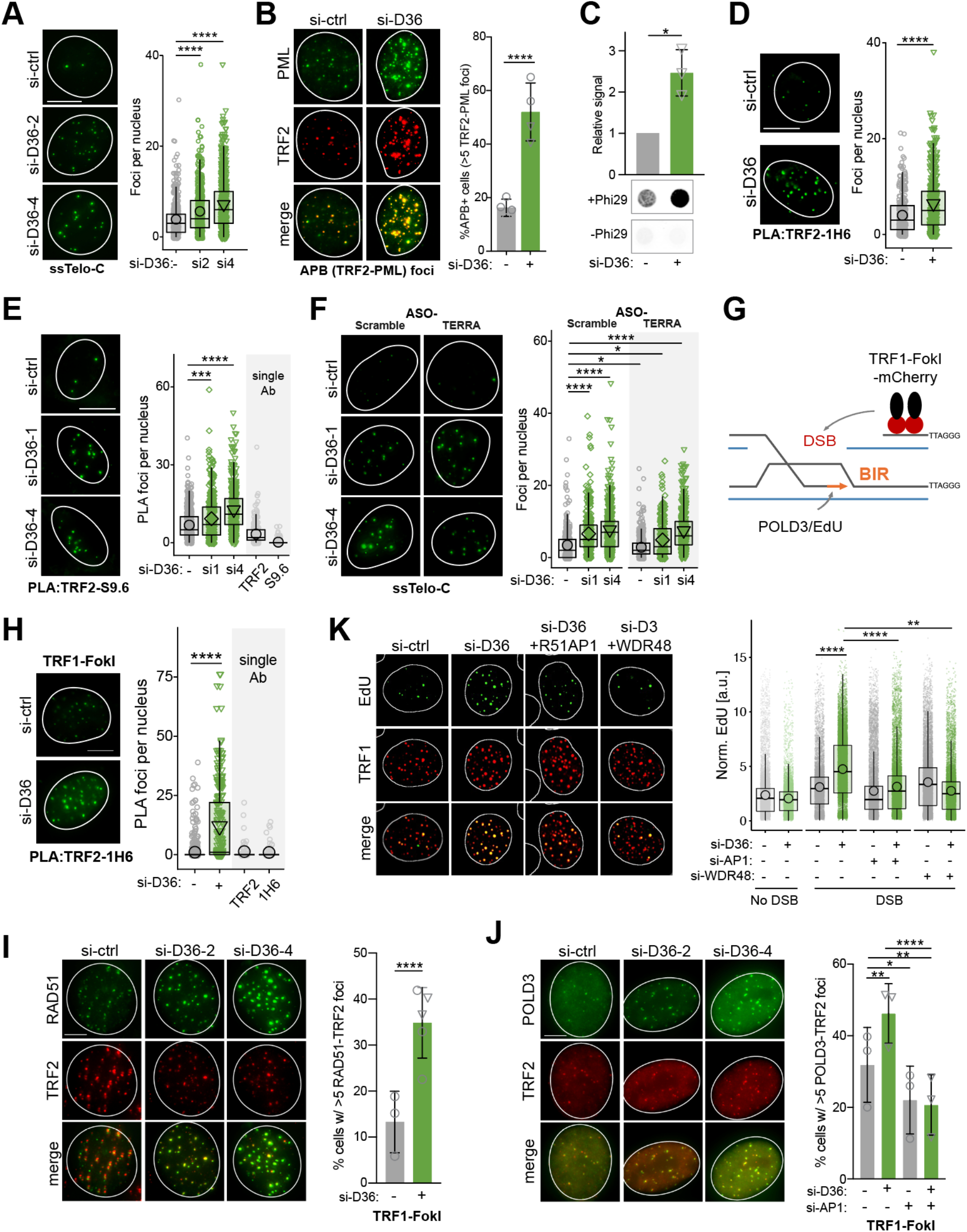
DHX36 depletion elevates ALT. **(A)** Quantitative IF using ssTelo-FISH in U2OS cells transfected with indicated siRNAs (n = 4, ≥ 75 cells per replicate), **(B)** Quantification of cells with >5 ALT-associated PML-bodies (APBs: TRF2-PML co-localization) in U2OS cells following transfection with indicated siRNAs (n=4, a min of 75 cells per replicate). **(C)** Rolling circle assay to quantify c-circles in U2OS cells transfected with the indicated siRNAs, bar graphs depict mean and S.D., symbols represent biological replicates (n = 3), P values based on unpaired T-test with Welch’s correction. **(D)** PLA to quantify G4s at telomeres using antibodies for G4s (1H6) and TRF2 (n = 2, ≥ 125 cells per replicate). **(E)** PLA to quantify R-loops at telomeres using S9.6 and TRF2 antibodies (n = 2, ≥ 100 cells per replicate). PLA with single antibody (TRF2, S9.6) served as control. **(F)** Depletion of TERRA with ASO transfection in cells transfected with the indicated siRNAs (n = 2, ≥ 100 cells per replicate), P values represent BH-adjusted pairwise comparisons from the fitted mixed-effects model. **(G)** Schematic of telomeric DSB induction via an mCherry-tagged TRF1-Fok1 fusion protein. **(H)** Analysis of telomeric G4s following the induction of TRF1-FokI-mCherry expression with Dox using PLA with G4 (1H6) and TRF2 antibodies (n = 2, ≥ 100 cells per replicate). **(I)** Analysis of RAD51 foci at telomeres following TRF1-FokI induction, symbols represent biological replicates (n = 3/5, ≥ 75 cells per replicate, P values based on a binomial mixed-effects model with biological replicate as a random effect, followed by Dunnett-adjusted comparisons to control. **(J)** Analysis of POLD3 foci at telomeres following TRF1-FokI induction, symbols represent biological replicates (n = 3), ≥ 40 cells per sample, mean and SD are indicated. **(K)** BIR activity following TRF1-Fok1 induced DSBs, box plots depict EdU mean intensities at TRF1 foci in G2/M cells from two independent biological replicates, ≥ 100 cells per sample. Unless noted otherwise, P values are based on Wilcoxon rank sum test with BH correction, * P < 0.05; ** P < 0.01; *** P< 0.001; **** P < 0.0001.

Telomeric DNA G4s are promoted by TERRA R-loop formation, and G4 formation can in turn stabilize R-loops (*9, 11*). Consistent with this, we found that DHX36 depletion elevated telomeric R-loops using PLA for S9.6 and TRF2, and that the depletion of SETX was epistatic with DHX36 depletion in ALT induction (**Fig. 4E, S5D**). We next asked whether TERRA were required for DHX36 loss-induced ALT by depleting TERRA RNA using a TERRA-targeting antisense oligonucleotide (ASO) or a scrambled control ASO (*37*). ASO treatment strongly reduced TERRA levels, and TERRA depletion reduced ALT in control cells consistent with previous work (**Fig 4F, S5E**) (*37*). However, TERRA ASOs had little effect on DHX36-depletion induced ALT, as ssTelo signal remained significantly elevated (**Fig 4F**). These data indicate that DHX36 loss can stabilize G4-mediated ALT largely independent of TERRA RNA, consistent with our findings in the DR-GFP reporter system.

To more closely mimic our observations in the DR-GFP assay, we employed the TRF1-FokI system that drives ALT through the generation of programmed breaks at the telomere (*14*) (**Fig 4. G**). DHX36 depletion led to a robust induction of telomeric G4s upon FokI induction (**Fig. 4H**). In addition, telomeric DSBs elevated RAD51 at telomeres, which was further increased upon DHX36 depletion, demonstrating HR activation consistent with our observations following IR in DHX36 depleted cells (**Fig. 4H, Fig. S2F**). Next, we assessed ALT activity downstream of template strand invasion by measuring DSB-induced accumulation of the BIR polymerase subunit POLD3, as well as telomeric EdU incorporation in G2/M cells (**Fig. 4G**). DHX36 loss enhanced both BIR hallmarks in a manner that was dependent on RAD51AP1 and WDR48 (**Fig. 4I, K, Fig. S5E**). Lastly, given that R-loop-driven ALT is RAD52-independent and required for the generation of extrachromosomal telomeric DNA species (*11, 13*), we examined the interplay RAD52 and DHX36. While DHX36 depletion led to elevated RAD52 localization at telomeres following FokI induction (**Fig. S5F**), RAD52 was dispensable for the DHX36 loss-induced increase in ALT activity (**Fig. S5G**). Together, these data establish that elevated G4s resulting from DHX36 loss can promote BIR in a RAD52-independent, RAD51AP1-dependent manner.

## Discussion

Here, we uncover DNA G4 dynamics as a central effector of homology search in somatic cells. Contrasting the prevailing view of DNA G4s as drivers of genome instability (*9, 38*), our findings show that stabilization of G4 DNA can promote HR-mediated genome maintenance by providing a substrate for template strand invasion. Open template DNA structures have a substantial kinetic advantage over recombinase- and ATP-dependent template strand unwinding (*6, 7*). G4s, thus, present a compelling means to ensure timely homology sampling, particularly in the context of the recently proposed, loop extrusion-mediated long-range scanning of RAD51 filaments against the sister chromatid (*3, 4*). While pre-existing G4s would provide suitable entry points, we favor a model wherein template scanning is coupled to G4 formation. Consistent with this, loop extrusion was recently shown to promote negative DNA supercoiling, which can in turn facilitate G4 formation (*39, 40*).

Depletion of DHX36, a G4 helicase linked to replication-associated G4 resolution, promoted HR via RAD51AP1/WDR48-mediated strand invasion, which can partially bypass the requirement for BRCA1 during HR. DHX36 depletion further restored RAD51 foci in BRCA1-deficient cells, pointing to additional G4 functions during RAD51 filament formation. Supporting a role for DNA secondary structures in the latter, both R loop formation at broken ends, as well as G4-mediated replisome stalling, have been linked to enhanced end resection (*33, 41, 42*).

While our findings indicate that G4s and R loops cooperate to facilitate HR in transcribed genomic regions to ensure their accurate repair (*43*), transcription is not a prerequisite for G4 formation (*9, 11*). Consistent with this, G4 stabilization upon DHX36 loss was able to facilitate HR in the absence of transcription. Transcription-independent, error-free repair may be particularly relevant in stem and progenitor cells where most tissue-specific genes are silenced, yet their mutagenesis would have detrimental effects for subsequent cell generations.

Contrasting their HR-promoting role, both G4s and R loops have been linked to replication stress-associated DNA damage (*9, 31, 38, 44*). Of note, loss of DHX36 or SETX was previously found to increase genome instability, suggesting that balanced secondary structure turnover is essential to ensure optimal genome maintenance (*31, 45*). Moreover, we cannot rule out that DHX36’s role in regulating mRNA translation contributes to its overall impact on genome maintenance (*46*). While ∼1% of the genome is predicted to have G4-forming potential (*12*), G4 structures are particulalry enriched at telomeric DNA due to the G-rich nature of the TTAGGG repeat, where they affect telomere maintenance (*9, 11, 38, 47*). Consistent with DHX36 function in HR, we observed increased ALT in response to DHX36 depletion, in a manner resembling the elevated ALT activity shown after treatment with G4 stabilizing compounds (*11, 48, 49*). In both cases, ALT activation was reminiscent of toxic hyper-ALT phenotypes elicited by hyperactivation of BIR, further underscoring the need for regulated G4 dynamics to prevent excessive genome instability (*49, 50*).

Our findings have direct implications for the targeting of DNA G4s in cancer therapy (*11, 38*). In the context of ALT, we provide additional support for the potential of G4 stabilizers as a means to enhance genome instability in ALT^+^ cancer patients (*49*). In the context of BRCA1 deficiency, our findings suggest caution regarding a potential restoration of HR and desensitization to HR-D-specific drugs, such as PARP inhibitors (*20, 21*). Consistent with this, G4 stabilizers showed no benefit in HR-D patients with BRCA1 mutations in a recent phase I trial in patients with solid tumors (*51*). In contrast, G4 stabilizing drugs were found to be cytotoxic in cells lacking BRCA1, BRCA2 or RAD51 in vitro (*52*), likely reflecting dose dependence and/or differences between global G4 stabilization and more selective G4 modulation upon DHX36 inactivation (*22*). It will be interesting to determine if additional DNA helicases and/or DNA/RNA secondary structures can similarly overcome HR-D upon BRCA1 loss, and whether this extends to other causes of HR-D. Altogether, our work establishes G4 stabilization as a driver of efficient homology search, and underlines the need to consider both adverse and genome-protective effects of DNA secondary structures.

## Acknowledgements

We thank T. Day, D. Durocher, R. Greenberg, E. Lazzerini-Denchi, G. Legube, A. Marin-Gonzalez, R.J. O’Sullivan, J. Vidigal, T.L. Wang and L. Zou for helpful discussions, critical reading, reagents and data sharing. We thank R. Chari of the NCI Genome Modification Core for preparation of the CRISPR library, Hao Zhang and Joe Margolick of the Flow Cytometry Cell Sorting Core Facility at Bloomberg School of Public Health, Johns Hopkins University for FACS sorting, the University of Michigan Vector Core for lentiviral library generation, the Center for Cancer Research (CCR) Genomics Core in Bethesda, Maryland, for help with high-throughput sequencing. Computations and data storage were supported by the Advanced Research Computing at Hopkins (ARCH) core facility (rockfish.jhu.edu) and the computational resources of the NIH HPC Biowulf cluster (https://hpc.nih.gov). The contributions of the NIH author(s) are considered Works of the United States Government. The findings and conclusions presented in this paper are those of the author(s) and do not necessarily reflect the views of the NIH and the U.S. Department of Health and Human Services.

## Authors contributions

Conceptualization: BLT, PB, SK, THS, PO; Methodology: BLT, PB, SK, SS, PJB, THS, PO; Validation: BLT, PB, SK, SS, YHL, THS, PO; Formal analysis: BLT, PB, SK, SS, YHL, TN, PJB, THS, PO; Investigation: BLT, PB, SK, SS, YHL, TN, VR, XZ; Resources: PJB; Data Curation: BLT, SK, SS, THS, PO; Writing – Original Draft: THS, PO; Writing – Review & Editing: BLT, PB, SK, SS, PBJ; Visualization: BLT, SK, SS, THS, PO; Supervision: BLT, PB, PJB, THS, PO; Project administration: PJB, THS, PO; Funding acquisition: PJB, THS, PO.

## Data, code, and materials availability

All data are available in the main text or the supplementary materials. All material generated in this study (including but not limited to oligonucleotides, plasmids and cell lines) are available on request. CRISPR/Cas9 screen NGS data generated for this study will be deposited at the Gene Expression Omnibus (GSE334161). Custom code used in this study will be deposited on GitHub.

## Materials and Methods

See **Table S1** for a list of antibodies, plasmids, chemicals, cell lines and software used in this study.

### Cell culture and treatments

HEK293T cells, U2OS osteosarcoma cells (ATCC) and their derivatives, LM216J, and LM216T cells were grown in Dulbecco’s modified eagle medium (DMEM, Invitrogen) supplemented with 10% FBS (Gemini) and maintained at 37 °C with 5% CO_2_. For U2OS-Tet-DR-GFP cells media was supplemented with 800 μg/ml G418, for RPE1-hTERT *TP53-KO, BRCA1-KO* cells, with 1% Glutamax (ThermoFisher) and 1% Penicillin/Streptomycin (ThermoFisher). UWB1.289 BRCA1^Δ^ and UWB1.289 BRCA1-reconstituted ovarian carcinoma cells were cultured in 1:1 RPMI-1640 and 50% Mammary Epithelial Growth Medium from (MEGM: MEBM basal medium and SingleQuot additives, Lonza) with 3% FBS. All cell lines were regularly tested for mycoplasma using Mycoplasma PCR detection kit (Abcam). 293T-DR-GFP and 293T-DR-GFP-Templ^G4Δ^ cells were generated by co-transfecting HEK293T cells with SpCas9 and an AAVS1-targeting sgRNA, and a template G4-mutant or wildtype pAAVS1-DR-GFP donor plasmid (see below for template G4 mutagenesis). Cells were selected with 250 µg/mL hygromycin for 10 days for stable transgene knock-in. For transient knockdown, cells were transfected with 25 nM siRNAs using DF-1 reagent following the manufacturer’s instructions (Dharmacon) and analyzed at 48 - 96 h post transfection. For experiments with multiple siRNA combinations, control siRNA was added as needed to produce equal siRNA concentrations. For stable knockdown, lentiviral infection of shRNA-expressing or LKO.1 empty vectors was carried out by spin infection (2500 rpm, 90 min, Eppendorf 5810R centrifuge) with 8 µg/ml polybrene (Sigma). Cells were incubated overnight prior to virus removal and selection with puromycin (Invitrogen, 2 µg/ml). See **Table S2** for siRNA and shRNA sequence information. Where indicated, cells were treated with IR (4GY X-ray, Xstrahl Biological Irradiator), Camptothecin (Selleckchem, 100 nM, 1 h), Olaparib (Selleckchem, 1.25 – 10 µM), Pyridostatin (Sigma, 2 µM), Phenylpyrrolocytosine (medchemexpress, 100 µM), Doxycycline (ThermoFisher Scientific, 1 – 5 µg/mL), Dimethyl sulfoxide (DMSO, Sigma) was used as vehicle control.

### HR reporter assays

(i) TRI-DR-U2OS cells carrying stably integrated DR-GFP reporter and Dox-inducible I-SceI transgenes were treated with 2.5 µg/mL Doxycycline for 48h and analyzed by flow cytometry for GFP expression as a proxy for HR efficiency (*17*). (ii) U2OS-Tet-DR-GFP cells were analyzed as described (*7*). In brief, cells were reverse transfected with siRNA using Lipofectamine® RNAiMAX Reagent (ThermoFisher). 24hrs after siRNA transfection, cells were transfected with I-SceI-T2A-mCherry (gift from Lee Zou) using Lipofectamine 3000 (ThermoFisher) according to the manufacturer’s instructions. Cells were reseeded and treated with 1 μg/mL Dox or vehicle (DMSO) for 62 h. A fraction of cells were subjected to FACS analysis to determine transfection efficiency (mCherry^+^ cells). Genomic DNA was prepared from the remaining cells using the DNeasy Blood and Tissue Kits for DNA Isolation kit (Qiagen). 75 ng of DNA was analyzed by qPCR for repaired DNA template (SceI-GFP) and a control locus (36B4), see **Table S2** for primer sequences. HR efficiency was calculated as the fraction of repaired template over control, normalized to transfection efficiency. (iii) 293T-DR-GFP and 293T-DR-GFP-Templ^G4Δ^ cells were transiently transfected with I-SceI-T2A-mCherry and analyzed 48 h thereafter by flow cytometry for HR efficiency, determined as % GFP^+^ cells divided transfection efficiency (% mCherry^+^). Where indicated, siRNA transfection was performed 24 h prior to I-SceI transfection.

### DR-GFP G4 mutagenesis

A pAAVS1-DR-GFP plasmid with synonymous mutations in predicted G4 sequences in the HR template region (position 9359 – 11609) was generated by Genscript, referred to as pAAVS1-DR-GFP-Templ^G4Δ^. Mutagenesis deleted all predicted G4s in the HR homology region, and several high scoring G4 sequences within close proximity, while maintaining 97% of the sequence homology between the DSB site and the template DNA. Imperfection-tolerant G-quadruplex prediction was performed using pqsfinder to validate disruption of predicted G4 motifs (*53*).

### Immunostaining and quantitative image-based cytometry (QIBC)

U2OS or TRI-DR-GFP cells were seeded and transfected with siRNAs as described above. 24 h after transfection, cells were reseeded in glass-bottom 96-well plates (Cellvis P96-1.5H-N). For IR-treated samples, cells were pulse-labeled with 10 μM EdU for 30 min immediately prior to harvest. For CPT recovery kinetics, EdU was pulse-labeled for 30 min prior to CPT treatment. For DSB induction at ALT telomeres, U2OS cells expressing TRF1-FokI-mCherry were transfected with siRNA as described and 48 hours later, treated with up to 3 µg/mL doxycycline for 16-24 hours and then 0.6 ug/ml 4-OHT (Sigma, 1μM) with or without Shield-1 (Takara Clontech, 1μM) for 2 hours. In selected experiments, cells were transfected with ASOs 48 h after the siRNA transfection and fixed for analysis 24 h later. For immunostaining, cells were pre-extracted on ice for 5 min using PBS-T extraction buffer containing 0.5% Triton X-100 (Sigma) in 1× PBS supplemented with protease and phosphatase inhibitor cocktail (Cell Signaling Technology), followed by fixation with 4% paraformaldehyde (Electron Microscopy Sciences) for 20 min at room temperature (RT). Cells were subsequently permeabilized with 0.5% Triton X-100 in PBS for 10 min. For G4 and S9.6 IF, cells were treated with RNAse A (Qiagen) (1 μg/ml) at 37°C for 30 min in PBS. EdU incorporation was detected using the Click-iT EdU Imaging Kit (ThermoFisher) according to the manufacturer’s instructions. Cells were blocked with 3% BSA (Sigma) in PBS for 1h at RT. Primary antibodies were diluted in PBS containing 2% BSA and incubated overnight at 4°C. Following washes with PBS 0.01% Tween-20 (PBS-T), cells were incubated with Alexa Fluor-conjugated secondary antibodies (ThermoFisher) for 1h at RT. For BIR, pre-extraction was omitted to retain TRF1-FokI-mCherry signal. Nuclear DNA was counterstained with DAPI (0.5 μg/mL, Invitrogen) for 5 min. Cells were mounted with Fluoromount-G (Sigma) and cured overnight at 4°C prior to imaging.

#### Image acquisition and analysis

Images were acquired using a Yokogawa SoRa spinning-disk confocal microscope equipped with a 20× objective and 2.8× magnification using full-sensor acquisition settings. Images were analyzed using custom Python-based analysis pipelines. Briefly, nuclear segmentation was performed using DAPI images with the Cellpose v3.0.8 built-in nuclei model. Partial nuclei and objects smaller than 500 pixels were excluded to minimize cell debris and segmentation artifacts. Nuclear foci were detected using combined LoG/DoG-based detection, merged to remove duplicate detections, and filtered by area and intensity, with foci required to exceed 1.5× the mean nuclear background intensity. Per-nucleus mean and integrated fluorescence intensities, EdU signal, and foci counts were quantified. At least 250 cells were analyzed per condition. For telomeric EdU (BIR) quantification, G2 cells were identified by QIBC cell-cycle profiling and EdU mean fluorescence intensity at TRF1 foci in G2 cells was assessed following normalization to the corresponding mean nuclear EdU intensity. At least 100 cells were analyzed per condition. For a subset of ALT assays, images were obtained using a Zeiss Axio observer and foci per nuclei were counted using Gen5 Software (Agilent BioTek). Statistical analysis was performed in R v4.5.1 using rstatix, and plots were generated with ggplot2. Box plots with jittered single-cell values show the interquartile range, median, mean (represented as large symbol), and whiskers extending to 1.5× IQR.

### Cell cycle analysis

Cell-cycle phases were defined based on integrated DAPI intensity and EdU incorporation. EdU-positive cells were identified for each experiment using the log-scaled EdU intensity distribution, where the cutoff was set between the two major populations. G1 cells displayed ∼2N DNA content with low/no EdU signal, S-phase cells showed positive EdU incorporation with intermediate DNA content, and G2 cells displayed ∼4N DNA content with low/no EdU signal.

### Proximity ligation assays

Proximity ligation assays were performed using the Duolink In Situ PLA kit (Sigma-Aldrich) according to the manufacturer’s instructions. 48h post siRNA transfection cells were seeded in the 18-chamber wells (Cellvis). Briefly, cells were fixed and permeabilized as in the QIBC protocol. Following permeabilization, samples were treated with RNase A (1 μg/ml) for 30 min at 37°C prior to blocking to remove RNA-dependent background signals. Cells were then blocked using Duolink blocking solution, incubated with FLAG-BG4 scFv fragment, then with anti-Flag antibody and/or rabbit α-RAD51 antibody for 2 h at 37°C. Next, samples were incubated with In situ PLA anti rabbit PLUS (Sigma) and In situ PLA anti mouse MINUS (Sigma) probes for 1 h at 37°C. Samples were incubated with ligation buffer and ligase for 30 min at 37°C, followed by incubation for 90 min at 37°C with phi28 polymerase for signal amplification. Reactions were terminated, nuclei were counterstained with DAPI (0.5 μg/ml for 5 min), coverslips were mounted in Fluoromount-G and cured overnight at 4°C. Imaging and quantitative analysis were performed as described for QIBC, with PLA foci quantified on a per-nucleus basis following segmentation.

### Native telomere FISH

Cells grown on coverslips were fixed in 2% paraformaldehyde for 5 min at RT, incubated with 500 µg/mL RNaseA (Millipore) in blocking solution (1 mg/mL BSA, 3% goat serum, 0.1% Triton X-100, 1 mM EDTA in PBS) for 1 h at 37°C. Following dehydration in ethanol (70%, 90%, and 100%) the TRITC-OO[TTAGGG]3-labeled PNA probe (PNA Bio) was added in hybridization buffer (70% formamide, 1 mg/mL blocking reagent (Roche), 10 mM Tris-HCl, pH 7.2) for 1 h at RT. Coverslips were washed 3 times with formamide washing buffer (70% formamide, 10 mM Tris-HCl, pH 7.2) and stained with DAPI. Imaging was carried out on a Lionheart automated microscope (Agilent BioTek) and nuclear signal quantified using Gen5 software (Agilent BioTekI). 2-3 independent experiments were performed unless indicated otherwise. For each experiment, a minimum of 50 of nuclei was imaged.

### C-circle assay

Genomic DNA was extracted from cells using DNeasy Blood and Tissue gDNA extraction kit (Qiagen), and DNA concentrations were quantified using Qubit dsDNA HS assay kit (ThermoFisher). Genomic DNA was used as the template for rolling circle amplification. Amplified products were then dot-blotted on nylon membrane, UV crosslinked and hybridized with TelC probe (5′-IRD800-CCCTAACCCTAACCCTAA-3′) from IDT. Membranes were scanned using LI-COR - ODYSSEY CLX and analyzed using Image J.

### Cellular extract preparation and immunoblotting

Cells were lysed in RIPA lysis buffer (25 mM Tris-HCl, pH 7.5; 150 mM NaCl; 2 mM EDTA; 1% NP-40; 1% Na-deoxycholate; 0.1% SDS) supplemented with complete^™^ EDTA-free protease inhibitor cocktail (Roche). Lysates were sonicated, centrifuged, diluted with 5x sample buffer (312.5 mM Tris-HCl, pH 6.8; 10% SDS; 50% glycerol; 12.5% β-mercaptoethanol; 0.05% bromophenol blue) and heated for 10 min at 95 °C. Lysates of equal protein amount based on BSA assay (Bio-Rad) were separated by SDS-PAGE and transferred onto nitrocellulose or PVDF membrane (Bio-Rad) at 100 V for 1 hour. Membranes were blocked with 5% milk powder (Bio-Rad) and incubated with specific antibodies at 4°C overnight. HRP-conjugated secondary antibodies were used for signal detection by enhanced chemiluminescence (Advansta). Immunoblots were visualized using a Li-COR Odyssey CLx system (LI-COR Biotechnologies) or a ChemiDoc MP imaging system (Bio-Rad).

### Chromatin immunoprecipitation

3 × 10^6^ cells were crosslinked with 1% formaldehyde for 10 min at RT, and crosslinking was quenched with glycine (final concentration 125 mM). Cells were washed twice with PBS and resuspended in cytoplasmic extraction buffer (10 mM HEPES, pH 7.9; 10 mM KCl; 1 mM EGTA; 0.25% NP40; 1× protease/phosphatase inhibitor cocktail) using a volume equal to five times the pellet size followed by incubation on ice for 5–10 min. Nuclei were collected by centrifugation at 2700 × g for 5 min at 4°C and washed twice with cytoplasmic extraction buffer lacking NP40. Isolated nuclei were treated with RNase A (1 μg/ml) at 37°C for 30 min. Nuclear pellets were resuspended in cold nuclear extraction buffer (20 mM HEPES, pH 7.9; 420 mM NaCl; 20% glycerol; 1 mM EDTA; 1× protease inhibitor cocktail and phosphatase inhibitor cocktail) and incubated on ice for 10 min. Chromatin fractions were collected by centrifugation at 5000 × g for 5 min at 4°C. Chromatin was sonicated using a Diagenode Bioruptor water bath sonicator (medium setting; 30 s ON/30 s OFF for 20 cycles), and supernatants were precleared with protein A/G beads. Samples were incubated with FLAG-BG4 scFv fragment for 1 h at 37°C, then with anti-Flag antibody overnight at 4°C together with 80 μl protein A/G beads. Beads were washed twice each with low-salt buffer, high-salt buffer, lithium chloride buffer, and TE buffer. Immunoprecipitated chromatin was eluted and reverse crosslinked overnight at 65°C. DNA was purified using PCR purification kit (Qiagen) and analyzed by quantitative PCR, see **Table S2** for primer sequences.

### MTT cell viability assay

For MTT assays, cells were seeded at 3000 cells per well in 96-well plates and treated with the indicated doses of Olaparib for 72 h. After the end of drug treatment, cells were treated with 0.5 mg/mL MTT reagent (Sigma) for 2 h, washed and incubated with 100 µL DMSO for at least 20 min. Absorbance was measured at 570 nm using SpectraMax M5 (Molecular Devices), with SoftMax Pro 5.2 software (Molecular Devices).

### RNA Extraction and RT-PCR

Total RNA was extracted using the TRIzol^™^ reagent according to the manufacturer’s instructions (Invitrogen). cDNA was synthesized from 1 µg of total RNA using PrimeScript RT Master Mix (Takara), and expression of the indicated genes was analyzed by quantitative RT-PCR using the CFX Opus 96 Real-Time PCR System (Bio-Rad) (see **Table S2** for primer sequences).

### G4 enrichment analysis at DSB-associated Hi-C contacts

G4 enrichment at DSB-associated Hi-C contacts was analyzed using predicted G-quadruplex (G4) coordinates from the PQS Finder database (hg38) and KR-normalized Hi-C data from SRA NCBI PRJNA1214218 (*3, 53*). Contacts were classified as DSB-associated or non-DSB controls. Two independently sampled non-DSB sets matched in contact number to the DSB set were merged for downstream analysis, and random genomic regions were included as an additional background control. Contacts within genomic distances <10 kb and >2 Mb were filtered to avoid low confidence interactions with contact score <6, DSB:DSB and self-contacts were excluded. DSB contacts passing these filters were ranked by contact score, and the top 5% were selected across sample groups.

G4 enrichment was quantified as G4 density (predicted G4 motifs per kilobase) within a 4 kb window centered on each contact anchor or control region. Matched upstream and downstream flanking regions located 10 kb from the contact site were analyzed to assess local specificity. Flanking regions were shifted away from the contact anchor when necessary to avoid overlap with the central contact window. Statistical significance was evaluated using permutation test (10,000 iterations), including two-sample comparisons between groups and paired comparisons between contact sites and matched flanks. Analyses were performed in R v4.5.1 using data.table and visualized with ggplot2.

### CRISPR screens and analysis

#### Library generation

A pooled CRISPR/Cas9 knockout library was designed to targeted 374 genes related to RNA modifications, secondary RNA/DNA structures and post-transcriptional processing, containing approximately 10 sgRNA per gene. The library is comprised of 3820 sgRNA and 100 control sgRNAs assembled into lentiGuide-Puro (gift from Feng Zhang; Addgene 52963) or pUSEPR backbones (gift from Joana Vidigal) as described previously (*54*). Pooled Libraries were transduced at MOI < 0.3 such that at least 500 cells received each sgRNA. *BRCA1* KO *TP53* KO RPE1-*hTERT* and TRI-DRGFP cells were selected with 20 µg/mL and 2 µg/mL puromycin, respectively.

#### Screen designs

To screen for modulators of PARPi sensitivity, sgRNA-transduced RPE1 hTERT p53^-/-^ BRCA1^-/-^ Cas9 cells were cultured for 3 days post selection, followed by treatment with 3.96 µM of Olaparib (Selleckchem) or DMSO (Millipore Sigma, control arm). Cells were passaged every two days for a total of 10 passages. After Olaparib selection, cells were passaged once more and cell pellets collected for genomic DNA preparation. To screen for modulators of HR efficiency, TRI-DR-GFP cells were transduced with Cas9-encoding lentiviral vector lentiCas9-Blast (Addgene 52962) at a MOI ∼10 and infected cells were selected with 2 μg/ml of blasticidin (ThermoFisher) for 10 days, followed by library transduction as above. Cells were split and maintained under selection for 1 week and then grown in complete culture medium for 3 days, followed by DSB induction with 5 µg/mL Dox. DMSO treatment or cells collected pre-treatment served as control arms. GFP^+^ cells, reflecting cells that have undergone HR, were sorted after 72 hours using a MoFlo XDP sorter (Beckman Coulter) for genomic DNA preparation.

#### DNA isolation and library preparation

DNA was isolated using the DNeasy Blood & Tissue kit or the QIAamp DNA Blood Mini Kit (Qiagen) and concentrated using Zymo DNA Clean & Concentrator-5 kit (Zymo), following manufacturer’s protocols. Up to 20 µg of DNA was amplified in a two-step PCR reaction. The first step PCR was performed with up to 250 ng of DNA per reaction, for samples with more than 250 ng of DNA, multiple PCRs per sample were set up to amplify the entirety of DNA from that sample. PCR was run on a QuantStudio3 light cycler (ThermoFisher) or CFX Opus 96 real time PCR system (Biorad). PCRs were stopped two to three cycles after exponential increase. First step PCR products were cleaned and pooled using 1X Ampure XP beads (Beckman) Products were quantified by Qubit™ dsDNA HS Assay Kit (ThermoFisher). Second step PCRs were performed on the entirety of product, split into multiple PCRs with a maximum of 20 ng template for each. PCR products were pooled and cleaned using 1X Ampure XP beads (Beckman). Barcoded PCR products were purified on a non-denaturing 10% polyacrylamide gel. After gel purification, samples were cleaned on Zymo DNA Clean & Concentrator-5 kit (Zymo) and submitted for Tape Station analysis with DNA 1000 regular sensitivity (Agilent Technologies). The final samples were pooled at equimolar amounts and sequenced on an Illumina NextSeq 500/550-series instrument using NextSeq High Output chemistry v2.5. See **Table S2** for primer sequences).

#### Data analysis

Sequencing adaptors were removed using TrimGalore (www.trimgalore.com/) and read alignment to the library, read counting, normalization, QC analysis of the samples, and calculation of the sgRNA counts LFC was carried out using MAGeCK-MLE (maximum likelihood estimation) (*55*). The MLE algorithm was used to estimate gene effect (beta score) and sgRNA efficiency. To identify effects on HR or Olaparib sensitivity, the Beta score difference (Beta-diff) was calculated (Dox-DMSO in the case of HR and Olaparib-DMSO in the case of PARPi sensitivity). For visual comparison of the 2 datasets, the Z-scores of the Beta-diffs were calculated. Genes were plotted in R-studio and genes with an FDR<0.05 in at least one of the screens and a Z >1 or <-1 are highlighted in Fig 1A (2026.04.0+526, Posit Software). See **Table S1** for relevant software. Sequencing data will be available upon publication via the NCBI Gene Expression Omnibus.

