## Supplemental Figures and Tables for "G quadruplex DNA facilitates a pervasive path to homologous recombination"

**Supplementary Materials**

**Figures S1-S5**

**Tables S1, S2**

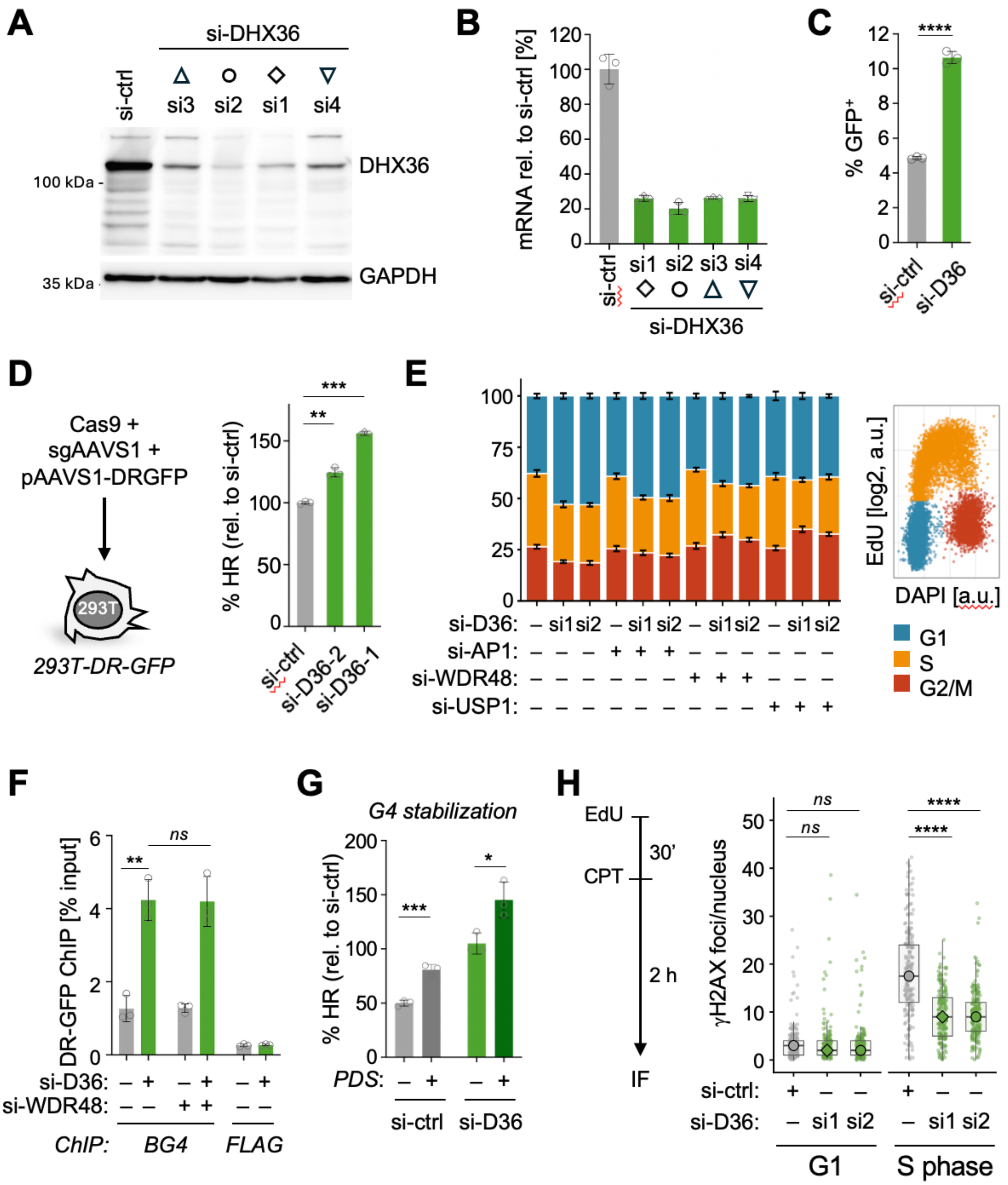

**Figure S1, related to Figure 1. (A)** Western blot for the indicated proteins in TRI-DR-GFP U2OS cells expressing the indicated siRNAs; symbols are used to define independent DHX36 siRNAs used in jitter and bar graphs throughout the manuscript. See Fig. S2B, C for HR efficiency upon transfection with siRNAs 3 and 4. **(B)** Quantitative PCR (qPCR) to assess DH36 knockdown efficiency in U2OS cells expressing the indicated siRNAs. mRNA levels were normalized to b-actin and are expressed relative to a control siRNA (si-ctrl). **(C)** Representative experiment showing the fraction of $GFP^{+}$ cells as a proxy for HR efficiency following I-SceI induction in TRI-DR-GFP cells expressing a control siRNA or DHX36-si2, used to calculate relative HR efficiency in Fig. 1C. **(D)** Left: Schematic for targeted insertion of the DR-GFP cassette into the AAVS1 safe harbor locus of 293T cells using CRISPR/Cas9 knock-in, resulting in 293T-DR-GFP cells. Right: HR efficiency in 293T-DR-GFP cells following transient transfection with I-SceI-T2A-mCherry. HR efficiency was measured as the fraction of $GFP^{+}$ cells normalized to transfection efficiency ($mCherry^{+}$) and is depicted relative to si-control (–). **(E)** Cell cycle profiles in cells expressing the indicated siRNA combinations. G1 (blue), S (orange) and G2/M phase (red) were defined based on DAPI and EdU intensities from IF images, a representative quantitative image-based cytometry (QIBC) profile analysis is shown. Bars depict mean and S.E.M. from 8 independent biological replicates. **(F)** G4 ChIP in TRI-DR-GFP cells expressing the indicated siRNA combinations in the absence of DSB induction. Enrichment at the DR-GFP template region is shown relative to input, FLAG IP served as a negative control. **(G)** HR efficiency as in Fig. 1C in TRI-DR-GFP cells expressing the indicated siRNAs in the presence or absence of PDS (2μM). **(H)** Representative IF analysis of γH2AX foci counts after 2 h recovery from CPT treatment, used to determine means shown in Fig. 1D. Box plots show mean intensities from at least 250 nuclei in G1 or S phase, P values are based on Wilcoxon test with BH correction. All bar graphs depict mean and S.D., symbols represent biological replicates. Unless noted otherwise, P values are based on Student's two-tailed t-test: * $P < 0.05$; ** $P < 0.01$; *** $P < 0.001$; **** $P < 0.0001$.

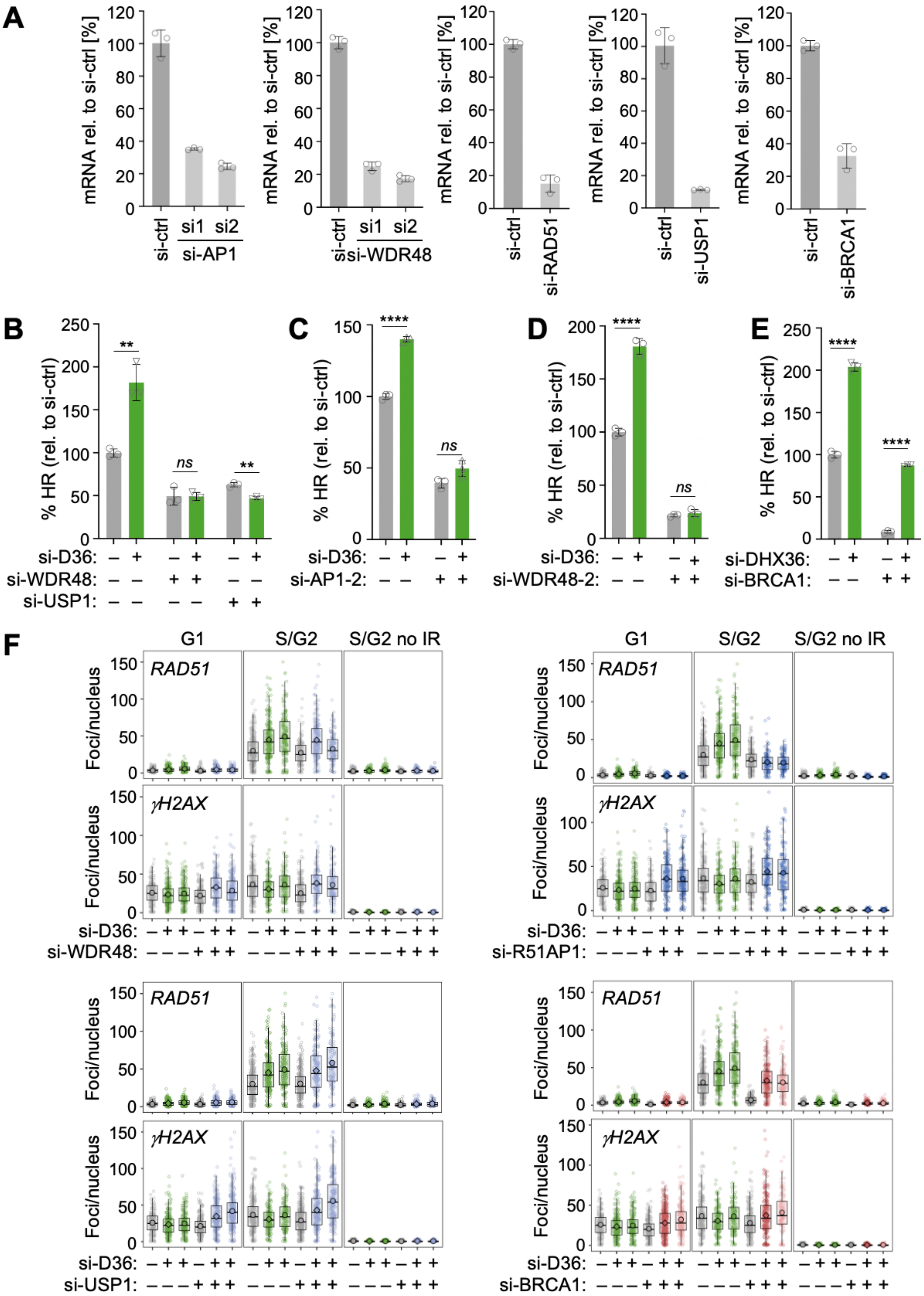
A
mRNA rel. to si-ctrl [%]
si-ctrl
si1
si2
si-AP1
si-WDR48
si-RAD51
si-USP1
si-BRCA1
B
% HR (rel. to si-ctrl)
**
ns
**
si-D36:
si-WDR48:
si-USP1:
C
****
ns
si-AP1-2:
D
****
ns
si-WDR48-2:
E
****
****
si-DHX36:
si-BRCA1:
F
G1
S/G2
S/G2 no IR
RAD51
γH2AX
Foci/nucleus
si-D36:
si-WDR48:
si-R51AP1:
si-USP1:
si-BRCA1:

**Figure S2, related to Figure 2. (A)** qPCR as in (Fig. S1B) to assess knockdown efficiencies for the indicated siRNAs in TRI-DR-GFP cells. **(B-E)** HR efficiencies as in Fig. 1C, using independent siRNAs for DHX36, RAD51AP1 (AP1) and WDR48. Distinct DHX36 siRNAs are depicted with distinct symbols, see Fig. S1A. **(F)** IF analysis of RAD51 and γH2AX foci counts in U2OS cells expressing the indicated siRNA combinations in G1 and S/G2 cells (see Fig. S1E). Box plots show mean intensities from at least 250 nuclei in G1 or S/G2 phase, P values are based on Wilcoxon test with BH correction. All bar graphs depict mean and S.D., symbols represent biological replicates. Unless noted otherwise, P values are based on student's two-tailed t test, symbols represent biological replicates. * P < 0.05; ** P < 0.01; *** P< 0.001; **** P < 0.0001.

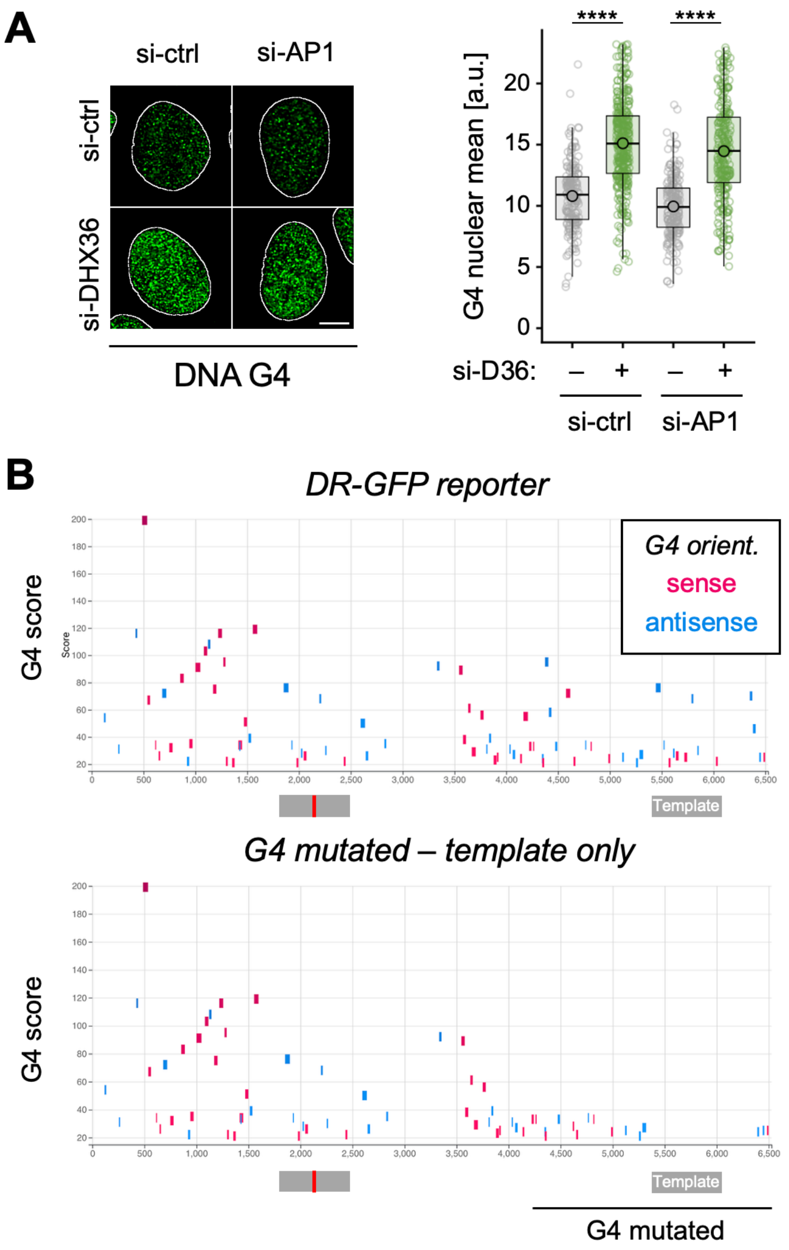

**Figure S3, related to Figure 2. (A)** Mean nuclear DNA G4 intensity in cells from Fig. 2D, analyzed as in Fig. 1B. Representative S phase nuclei are shown, scale bar: 5 μm, n > 250 nuclei. P values are based on Wilcoxon rank-sum test with BH correction, **** P < 0.0001. **(B)** G4 predictions based on pqs finder (see Methods) for the DR-GFP and $Templ^{G4\Delta}$ DR-GFP transgenes. Gray boxes depict regions of homology, the red line the I-SceI site. Black line demarks region wherein high-scoring G4s were mutated. All predicted G4s were mutated within the template.

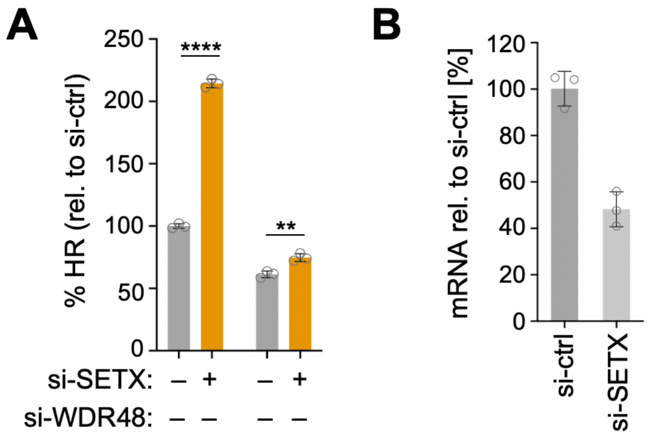

**Figure S4, related to Figure 3. (A)** HR efficiencies in TRI-DR-GFP cells as in Fig. 3C, using an independent siRNA for WDR48. **(B)** qPCR as in Fig. S2B to assess knockdown efficiency of SETX in TRI-DR-GFP cells. All bar graphs depict mean and S.D., symbols represent biological replicates. P values are based on Student's two-tailed t-test: * P < 0.05; ** P < 0.01; *** P< 0.001; **** P < 0.0001.

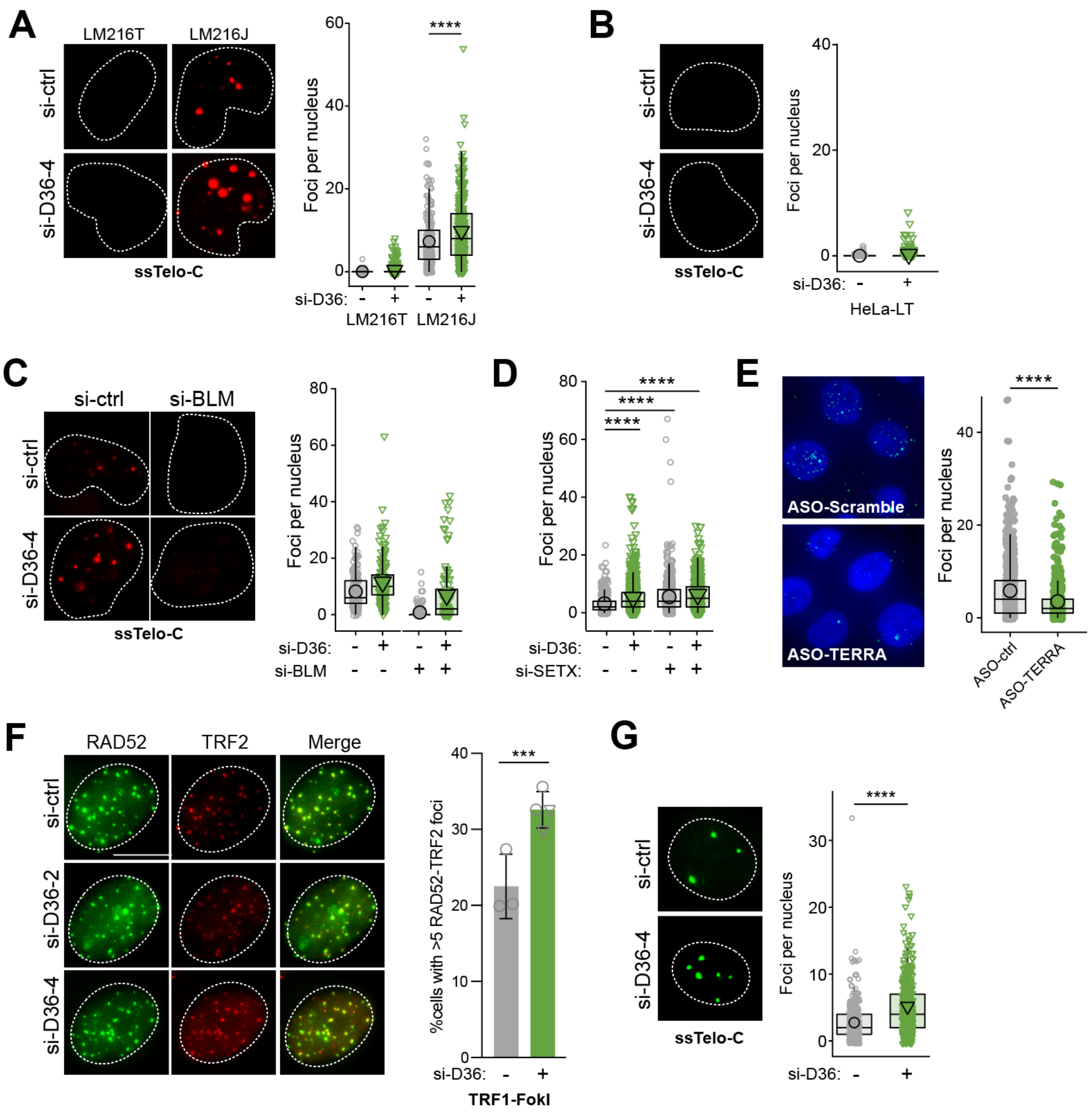

**Figure S5, related to Figure 4. (A)** Quantitative IF using ssTelo-FISH in LM216J (ALT⁺) and LM216T (hTERT⁺) cells transfected with indicated siRNAs (n=2,4, ≥ 50 cells per replicate). **(B)** ssTelo-FISH in HeLa-LT cells transfected with indicated siRNAs (n = 2, ≥ 75 cells per replicate. (C) ssTelo-FISH in U2OS cells transfected with the indicated siRNAs (n = 2, ≥ 40 cells per replicate). **(D)** ssTelo-FISH in U2OS cells transfected with indicated siRNAs (n = 3, ≥ 100 cells per replicate). **(E)** FISH analysis of TERRA foci following transfection with scramble or TERRA targeting ASOs. **(F)** Analysis of RAD52 foci at telomeres following TRF1-FokI induction (n = 3,4), P values are based on a binomial mixed-effects model with biological replicate as a random effect, followed by Dunnett-adjusted comparisons to control. **(G)** ssTelo-FISH in U2OS RAD52 KO cells transfected with indicated siRNAs (n = 2, ≥ 80 cells per replicate). Unless noted otherwise, P values based on Wilcoxon rank sum test with BH correction, * P < 0.05; ** P < 0.01; *** P< 0.001; **** P < 0.0001.

### Table S1: Reagents and Tools

| Primary Antibodies | | | | |
|---|---|---|---|---|
| **Antibody** | **Vendor** | **Application** | **Dilution** | **Identifier (Cat#, RRID)** |
| G4 (BG4) | Sigma, MABE917 | IF<br>PLA<br>IF | 1:5,000<br>1:300<br>1:300 | RRID:AB_2750936 |
| G4 (1H6) | Sigma, MABE1126 | PLA | 1:800 | RRID:AB_2924428 |
| R-loop (S9.6) | Sigma, MABE1095 | PLA | 1:500 | RRID:AB_2861387 |
| PML | Santa Cruz, sc966 | IF | 1:500 | RRID:AB_628162 |
| BLM | Santa Cruz, sc365753 | Western | 1:1,000 | RRID:AB_10851630 |
| RAD51 | BioAcademia, 70-002 | IF | 1:20,000 | RRID:AB_1056187 |
| RAD51 | CST, 65653 | PLA<br>IF | 1:300<br>1:300 | RRID:AB_3718052 |
| RAD52 | Santa Cruz, sc365341 | IF | 1:250 | RRID:AB_10851346 |
| POLD3 | Abnova, H00010714-M01 | IF | 1:250 | RRID:AB_606803 |
| TRF2 | Novus, NB110-57130 | PLA<br>IF | 1:1,000<br>1:1,000 | RRID:AB_844199 |
| TRF2 | Invitrogen, MA1-41001 | IF | 1:1,000 | RRID:AB_2201326 |
| α-DHX36 | Abcam, ab226813 | Western | 1:3,000 | RRID:AB_2735719 |
| α-FLAG | Sigma, F1804 | IF<br>ChIP<br>PLA | 1:100<br>1:100<br>1:100 | RRID:AB_262044 |
| α-GAPDH | Santa Cruz, sc47724 | Western | 1:3,000 | RRID:AB_627678 |
| α-gH2AX | Millipore, JBW301 | IF | 1:500 | RRID:AB_2847865 |
| **Secondary Antibodies** | | | | |
| Goat anti-Mouse IgG (H+L) Alexa Fluor 488 | ThermoFisher, A11001 | IF | 1: 500 | |
| Goat anti-Rabbit IgG (H+L) Alexa Fluor 488 | ThermoFisher, A11008 | | 1:500 | |
| Goat anti-Rabbit IgG (H+L) Alexa Fluor 568 | ThermoFisher, A11011 | IF | 1:500 | |
| Goat anti-Mouse IgG (H+L) Alexa Fluor 647 | ThermoFisher, A21235 | | 1:500 | |
| Goat anti-Mouse IgG-HRP | Jackson ImmunoResearch | Western | 1:10,000 | |
| Goat anti-Rabbit IgG-HRP | Jackson ImmunoResearch | Western | 1:10,000 | |

| | | | |
|---|---|---|---|
| Goat Anti-Mouse IgG H&L (Alexa Fluor® 488) preadsorbed | Abcam, ab150117 | IF | 1:500 |
| Goat Anti-Rabbit IgG H&L (Alexa Fluor® 647) preadsorbed | Abcam, ab150083 | IF | 1:500 |
| Goat Anti-Mouse IgG H&L (Alexa Fluor® 647) preadsorbed | Abcam, ab150119 | IF | 1:500 |
| Goat Anti-Rabbit IgG H&L (Alexa Fluor® 488) preadsorbed | Abcam, ab150081 | IF | 1:500 |

| **Recombinant DNA** | |
|---|---|
| **Plasmid** | **Source** |
| pSANG10-3F-BG4 | Addgene 55756 |
| pX458-AAVS1-sg | Addgene, 194721 |
| pAAVS1-DR-GFP | Addgene, 113193 |
| pLKO.1-shDHX36-puro | Sigma, TRCN0000290002 |
| pLKO.1-puro | Addgene, 10878 |
| lentiCas9-Blast | Addgene, 52962 |
| I-SecI-T2A-mCherry | Gift from L, Zou, Duke University |

| **Chemicals and Reagents** | | |
|---|---|---|
| **Chemical** | **Vendor** | **Cat#** |
| Olaparib | Selleckchem | S1060 |
| Doxycycline | ThermoFisher Scientific | 24390-14-5 |
| Lipofectamine RNAiMAX | ThermoFisher Scientific | 13778150 |
| Qubit dsDNA HS assay kit | ThermoFisher Scientific | Q32851 |
| Triton X-100 | ThermoFisher Scientific<br>Sigma | BP151-500<br>282103 |
| paraformaldehyde | Electron Microscopy Sciences | 15713 |
| RNAse A | Qiagen | 19101 |
| RNase H1 | NEB | M0297S |
| DAPI | Invitrogen | D1306 |
| Fluoromount-G Mounting Media | Sigma | F4680 |
| Protease/phosphatase inhibitor | Cell Signaling Technology | 5872 |
| complete™ EDTA-free protease inhibitor | Roche | 11873580001 |

| | | |
|---|---|---|
| MTT reagent | Sigma | 475989 |
| Trizol | Invitrogen | 15596026 |
| PrimeScript RT Master Mix | Takara | RR036A |
| Click-iT EdU Imaging Kit | ThermoFisher Scientific | C10634, C10633 |
| Duolink In Situ PLA kit | Sigma | DUO92101 |
| In situ PLA anti rabbit PLUS | Sigma | DUO92002 |
| In situ PLA anti mouse MINUS | Sigma | DUO92004 |
| Bovine Serum Albumin (BSA) | Sigma | A9647 |
| RNAse A | Qiagen | 19101 |
| Pyridostatin | Sigma | SML2690 |
| EdU (5-ethynyl-2'-deoxyuridine) | ThermoFisher Scientific | A10044 |
| PhcP | medchemexpress | HY-164421 |
| Camptothecin | Selleckchem | S1288 |
| Dimethyl sulfoxide | Sigma | 34869 |
| Puromycin | Invitrogen | 58-58-2 |
| Hygromycin B | ThermoFisher Scientific | 10687010 |
| Polybrene | Sigma | TR-1003 |
| Blasticidin | ThermoFisher Scientific | A1113903 |

| **Experimental models: Cell lines** | | |
|---|---|---|
| **Cell line** | **Source** | **Reference / Cat#** |
| U2OS | ATCC | Cat# HTB-96; RRID:CVCL |
| U2OS RAD52 KO | Gift from L. Zou, Duke University | *(11)* |
| U2OS-TRI-DR-GFP | | *(17)* |
| U2OS-Tet-DR-GFP | Gift from L, Zou, Duke University | *(7)* |
| U2OS mCherry-TRF1-FokI | Gift from R. Greenberg, University of Pennsylvania | *(14)* |
| LM216J | Gift from R. O'Sullivan, University of Pittsburgh | *(34)* (no CVCL) |
| LM216T | Gift from R. O'Sullivan, University of Pittsburgh | *(34)* (no CVCL) |
| HeLa-LT | Gift from E. Lazzerini-Denchi, NCI | *(34)*; CVCL_F0J8 |
| HEK293T | ATCC; gift from The Broad Institute, Cambridge | |
| UWB1.289 BRCA1$^{D}$ and UWB1.289 BRCA1-reconstituted OvCa cells | Gift from T.-L. Wang, Johns Hopkins University | *(56)* |
| RPE1-hTERT *TP53-KO*, *BRCA1-KO* | Gift from D. Durocher, University of Toronto | *(19)* |

| **Software and algorithms** | | |
|---|---|---|
| **Software** | **Vendor** | **Reference** |

| | | |
|---|---|---|
| Prism v11.0.1 | GraphPad | |
| R-studio version 2026.4.0.526 | Posit team (2026) | RStudio: Integrated Development Environment for R. Posit Software, PBC, Boston, MA. URL http://www.posit.co/ |
| R version X | R Core Team (2026) | R: A Language and Environment for Statistical Computing_. R Foundation for Statistical Computing, Vienna, Austria. doi:10.32614/R.manuals |
| Lionheart FX (Gen5) | Agilent BioTek (ABI) | |
| Zen 2.6 (blue edition) | Zeiss | |
| CellPose v3.0.8 | Open source | *(57)* |
| MAGeCK MLE | Open source | *(55)* |
| ChatGPT | Open AI (2026) | *ChatGPT (May 5 version)* [Large language model]. https://chat.openai.com/ |
| | | Portions of the code were developed with assistance from ChatGPT (OpenAI, 2026), a large language model. The authors reviewed and validated all outputs and take full responsibility for the final implementation and its correctness. |
| TrimGalore | Open source | www.trimgalore.com/ |

**Table S2: Oligonucleotide Sequences**

| shRNA | | | |
|---|---|---|---|
| **Name** | **Sequence** | **Source** | **Notes** |
| **shRNA** | | | |
| sh-DHX36-2 | 5'-CGACGAGAAGAACAAATTGTA-3' | Sigma | TRCN0000290002 |
| **siRNA** | | | |
| **Name** | **Sequence** | **Source** | **Notes** |
| si-DHX36-1 | 5'-GUAAAGACUUAAUGUGCAUGACUTG-3' | IDT | hs.Ri.DHX36.13.3 |
| si-DHX36-2 | 5'-CGACGAGAAGAACAAATTGTA-3' | Dharmacon | |
| si-DHX36-3 | 5'-TCCGCTGAGTGGGTTAGTAAA-3' | Dharmacon | |
| si-DHX36-4 | 5'-ACGCUUUGGAUAAACAAGAAGAATT-3' | IDT | hs.Ri.DHX36.13.1 |
| si-RAD51AP1 | 5'-GGAACUUCCAACAGUCACCACUAAT-3' | IDT | hs.Ri.RAD51AP1.13.3 |
| si-R51AP1-2 | 5'-CUGUUUUUCUAUCAGUUCGACAUGA-3' | IDT | hs.Ri.RAD51AP1.13.1 |
| si-WDR48 | 5'-CAAGGGAUUUUGCAUGUCAACAUTA-3' | IDT | hs.Ri.WDR48.13.1 |
| si-WDR48-2 | 5'-CCAAGGAUAAAGAACUAGUAGCATC-3' | IDT | hs.Ri.WDR48.13.3 |
| si-USP1 | 5'-AGUAUGAGGCAUUCUGAAGACUUTA-3' | IDT | hs.Ri.USP1.13.1 |
| si-BRCA1 | 5'-GUACGAGAUUUAGUCAACUUGUUGA-3' | Qiagen | SI02654575 |
| si-SETX | 5'-GCCAGAUCGUAUACAAUUAU-3' | Dharmacon | *(58)* |
| si-FANCJ | 5'-AGAAGAUAAACAGUCCACUUCAAAT-3' | IDT | hs.Ri.BRIP1.13.2 |
| si-control | ON-TARGET Plus | Dharmacon | |

| **guideRNA** | | | |
|---|---|---|---|
| **Name** | **Sequence** | **Source** | **Notes** |
| sgAAVS1 | 5’- GGGGCCACTAGGGACAGGAT-3’ | Addgene | Plasmid 113194 |
| **ASO** | | | |
| **Name** | **Sequence** | **Source** | **Notes** |
| ASO-Scrambled | AACACGTCTATACGC | Qiagen | *(37)* |
| ASO-TERRA | TAACCCTAACCCTAAC | Qiagen | *(37)* |
| **ssTelo probes** | | | |
| **Name** | **Sequence** | **Source** | **Notes** |
| TelG-FAM | CCCTAACCCTAACCCTAA | PNA Bio F1005 | *(34)* |
| TelG-Cy3 | TTAGGGTTAGGGTTAGGG | PNA Bio F1006 | *(34)* |
| **Primers (IDT)** | | | |
| **Name** | **Sequence** | **AT*** | **Application** |
| SceI-GFP F | 5'-GGGCGATGCCACCTACG-3' | 67 | Tet-DR-GFP *(7)* |
| SceI-GFP R | 5'-GGTGTTCTGCTGGTAGTGGTCG-3' | 67 | Tet-DR-GFP *(7)* |
| 36B4 F | 5'-CAGCAAGTGGGAAGGTGTAATCC-3' | 60 | Tet-DR-GFP *(7)* |
| 36B4 R | 5'-CCCATTCTATCATCAACGGGTACAA-3' | 60 | Tet-DR-GFP *(7)* |
| G4 ChIP F | 5’-GCCGCGACTCTAGATCATAA-3’ | 60 | ChIP |
| G4 ChIP R | 5’-CCACAACTAGAATGCAGTGAAA-3’ | 60 | ChIP |
| b-Actin RT F | 5’-TTCTACAATGAGCTGCGTGTGGCT-3’ | 60 | RT-PCR |
| b-Actin RT R | 5’-TCATCTTCTCGCGGTTGGCCT-3’ | 60 | RT-PCR |
| DHX36 F | 5'-GAGGCAGTGTTACTCTCCATAAG-3' | 60 | RT-PCR |
| DHX36 R | 5'-TATGTGGCTCAACGGGTAATC-3' | 60 | RT-PCR |
| RAD51AP1 F | 5'-TGCAGTGTAGCCAGTGATTATT-3' | 60 | RT-PCR |
| RAD51AP1 R | 5'-TGCTGTGCTGCAGCTTTA-3' | 60 | RT-PCR |
| WDR48 F | 5'-CTCACTGCTAGTCAGCTTCTTT-3' | 60 | RT-PCR |
| WDR48 R | 5-'GAGGAAGGGAGAAAGACCATTT-3' | 60 | RT-PCR |
| USP1 F | 5'-GAACAGCTCCAGGCTAGTTT-3' | 60 | RT-PCR |
| USP1 R | 5'-TGAGTTCCCTCAGTGTGTTAAG-3' | 60 | RT-PCR |
| BRCA1 F | 5'-ACCTACAACTCATGGAAGGTAAAG-3' | 60 | RT-PCR |
| BRCA1 R | 5'-CTTCAGCTCTGGGAAAGTATCG-3' | 60 | RT-PCR |
| FANCJ F | 5'-CCAGCTGGAGGCTAATCATATC-3' | 60 | RT-PCR |
| FANCJ R | 5'-CTGGAAGGTAGCACAGAGATTC-3' | 60 | RT-PCR |
| step 1 F primer pool (N = 1 to 8 random bp) | 5’-TCC CTA CAC GAC GCT CTT CCG ATC TNC TTG TGG AAA GGA CGA AAC ACC G-3’ | 60 | NGS library *(54)* |
| step 1 R lentiGuide-Puro | 5’-GTT CAG ACG TGT GCT CTT CCG ATC T**CT CTG CTG TCC CTG TAA TAA**-3’ | 60 | NGS library *(54)* |
| step 1 R, pUSEPR | 5’-GTT CAG ACG TGT GCT CTT CCG ATC T**TC TCT AGG CAC CGG TTC AAT**-3’ | 60 | NGS library *(54)* |

| | | | |
|---|---|---|---|
| step 2 'universal' F | 5’-AAT GAT ACG GCG ACCA CCG AGA TCT ACA 5’-C**TA GAT CGC** ACA CTC TTT CCC TAC ACG ACG CTC TTC CGA TC*T-3’ | 61 | NGS library *(54)* |
| step 2 'universal' R; X: unique bar code | 5’-CAA GCA GAA GAC GGC ATA CGA GAT **XXX XXX XX**GT GAC TGG AGT TCA GAC GTG TGC TCT TCC GAT C*T-3’ | 61 | NGS library *(54)* |
